# Cell-type specific transcriptional modulation by psilocybin induces sustained plasticity in mouse medial prefrontal cortex

**DOI:** 10.1101/2025.01.08.631940

**Authors:** Heike Schuler, Delong Zhou, Chloé Savignac, Vedrana Cvetkovska, Yiu-Chung Tse, Juliet Meccia, Joëlle Lopez, Ashot S. Harutyunyan, Jiannis Ragoussis, Danilo Bzdok, Rosemary C. Bagot

## Abstract

Despite enormous interest in psychedelics for psychiatric interventions, potential underlying biological mechanisms remain unclear. Here, we confirm that a single dose of psilocybin increases synaptic transmission in mouse medial prefrontal cortex. Using scRNA-sequencing, we identify cell-type specific mechanisms of sustained neuroplastic effects. We show that, 24h post-psilocybin, expression of plasticity-related genes is increased in excitatory neurons and that transcription in a type of deep layer near projecting neuron, L5/6 NP, is robustly altered. Analyzing receptor expression patterns reveals that this cell-type specificity does not align with 5-HT_2A_ expression but aligns with 5-HT_2C_ expression patterns. Further, multivariate analyses identify psilocybin-induced gene expression patterns in L5/6 NP neurons predict 5-HT_2C_, but not 5-HT_2A_, transcript levels. Pharmacologic manipulation with a 5-HT_2C_ antagonist attenuates the post-acute sustained effect of psilocybin on synaptic transmission, highlighting 5-HT_2C_ signaling and L5/6 NP neurons as key mediators of psychedelic drug action’s sustained neuroplastic effects in mPFC.

## Background

In a mounting mental health epidemic, claims that psilocybin, a psychedelic drug, provides lasting relief from a range of psychiatric disorders have fueled an explosion of interest^1–12^. Psilocybin, a serotonergic hallucinogenic compound from fungi, induces both acute and enduring effects^13^. Acutely, psilocybin induces hallucinations and altered states of consciousness^14^. Psilocybin also exerts sustained therapeutic effects in disorders including depression, anxiety and alcohol use disorder^1–3,5–7^. Psilocybin is rapidly metabolized into, psilocin, which ubiquitously binds serotonin receptors^15,16^.

While 5-HT_2A_ receptors mediate the hallucinogenic effects^14,17^, the mechanism of sustained effects is unclear and may involve serotonin receptors beyond 5-HT_2A_ ^18,19^. Understanding the lasting impact of psilocybin on diverse cell types is critical to safe clinical use.

The medial prefrontal cortex (mPFC) mediates cognition and mood, is implicated across psychiatric disorders^20,21^ and is a key locus of psilocybin’s therapeutic effects^19,22–26^. In humans, psilocybin regulates prefrontal cortical networks^22,27^. In rodents, psilocybin enduringly increases dendritic spines, excitatory neurotransmission, and BDNF-mTOR signaling potentially interacting with the BDNF receptor, TrkB^19,28,29^. Recent work suggests that plasticity in layer 5PT neurons may be particularly important for the potential therapeutic effects of psilocybin^30^. These phenomena are translationally relevant: treating human cortical iPSCs with psilocin also increases neurite complexity, excitatory synaptic transmission and neuroplasticity-associated gene expression^31^.

While recent findings suggest cell-type specificity, it is unclear how broadly psilocybin impacts neuronal and non-neuronal cell types in mPFC^30^. We used single-cell RNA sequencing (scRNAseq) to probe lasting transcriptional changes in diverse cell-types in the mouse mPFC after a single injection of psilocybin, at a timepoint when excitatory synaptic transmission is increased. Leveraging machine learning methods to predict treatment from individual cell-types and identify treatment-predictive gene modules, we find that psilocybin preferentially impacts a previously overlooked cell-type, L5/6 NP neurons. We show that L5/6 NP neurons enrich for the gene encoding 5-HT_2C_ receptors, *Htr2c,* and that psilocybin-associated gene module expression predicts *Htr2c* expression. Finally, pre-treatment with a 5-HT_2C_ antagonist attenuates increased synaptic transmission after psilocybin, confirming 5-HT_2C_ as a key regulator of psilocybin’s sustained neuroplastic effects.

## Results

### Psilocybin induces lasting increases in synaptic transmission in layer 5/6 mPFC neurons

We first established the dose-response curve for psilocybin’s acute effects on behavior. Consistent with published reports, 1mg/kg maximally increased head-twitch, a measure of 5-HT_2A_ signaling (Fig.1a,b; Table S1)^17^. Psilocybin may also induce sustained behavioral changes in rodents but effects can be variable^18,19,32–37^. In contrast, synaptic function provides a robust, reproducible, translationally-relevant assay of psilocybin’s sustained effects^18,19,30,31^. We first asked if a single injection of psilocybin enduringly increases synaptic transmission in infralimbic mPFC, similar to motor/cingulate cortex^19^ and hippocampus^18^. We injected mice with psilocybin (1mg/kg), prepared acute slices for patch-clamp recordings 24h later and recorded mEPSCs in neurons located in layers 2/3 or 5/6 (Fig. 1c). In layer 5/6 neurons, psilocybin increased mEPSC frequency and amplitude (Fig. 1d-f). In layer 2/3 neurons, psilocybin did not alter average frequency or amplitude indicating a degree of cell-type specificity.

**Figure 1.**
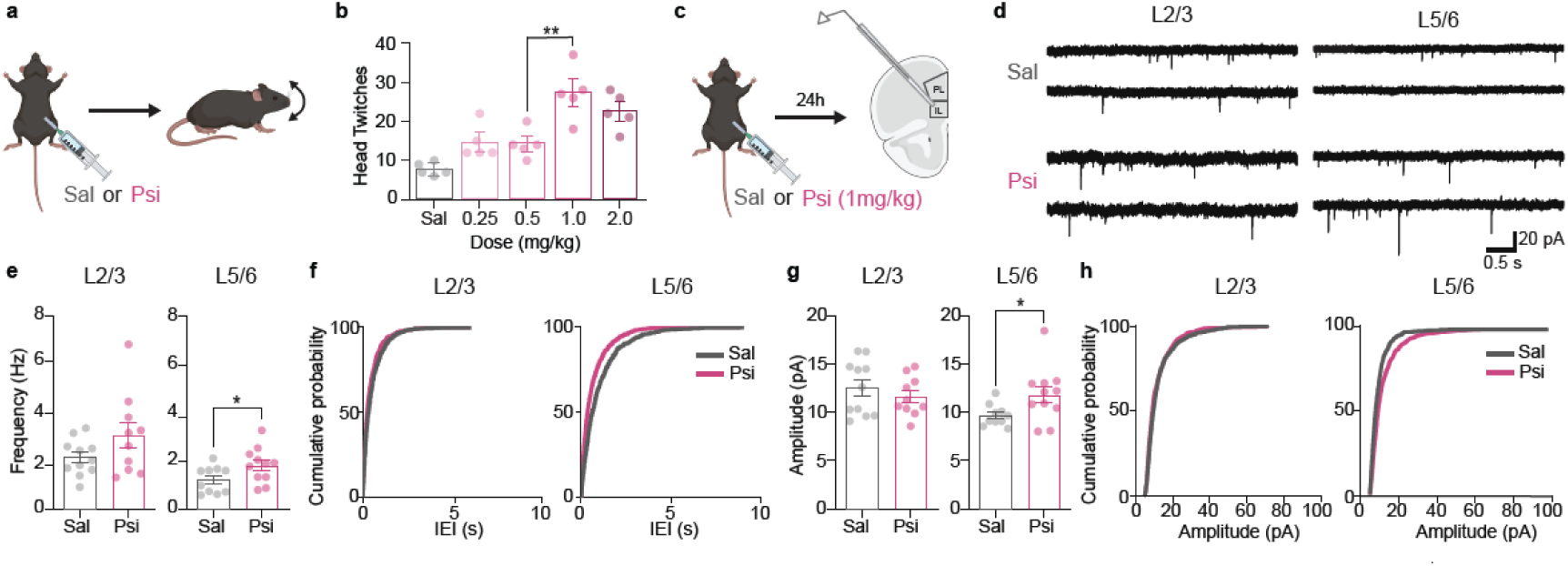
A single dose of psilocybin induces a sustained increase in synaptic transmission in medial prefrontal cortex. **a**. Schematic of head-twitch dose-response experiment. Mice were injected with psilocybin (0.25, 0.5, 1 or 2mg/kg) or saline and placed in a clean cage to observe head-twitch response for 30 minutes. **b.** Total head-twitch response (HTR) induced by psilocybin. HTR peaked at 1mg/kg and exceeded HTR induced by 0.5mg/kg (n=5; one-way ANOVA, p<0.0001, p=0.002, Sidak post-hoc test). **c.** Schematic of synaptic physiology experiment. Mice were injected with psilocybin (1mg/kg) or saline and placed in a clean cage for 30 mins then returned to their home-cage. 24h later brain slices containing the infralimbic cortex were prepared for patch-clamp recordings. **d.** Representative traces of mEPSC recorded from L2/3 and L5/6 neurons in the infralimbic cortex in brain slices from mice that received saline of psilocybin 24h prior to recording. **e.** Bar graphs of mEPSC frequency. Psilocybin selectively increased mEPSC frequency in L5/6 neurons (p<0.05), but not L2/3 neurons. Each circle denotes a cell (L2/3: n=11 cells from five mice for saline, and n=10 cells from six mice for psilocybin; L5/6: n= 10 cells from 5 mice for saline, and n= 11 cells from six mice for psilocybin). **f.** Cumulative distribution plots of mEPSC inter-event interval (IEI) for the same recordings. Psilocybin selectively decreased IEI in L5/6 neurons. **g,h**. Similar to **e, f** but for mEPSC amplitude. Psilocybin increased amplitude in L5/6, but not L2/3 neurons (p<0.05).

### Single cell RNA-Seq reveals psilocybin induces neuroplasticity gene programs

In ubiquitously binding 5-HTRs which are widely expressed, psilocin can impact many cell-types^15,17,38^. To examine the sustained effect of psilocybin in mPFC neuronal and non-neuronal cells, we used scRNA-seq. scRNAseq presents certain advantages over single nucleus RNAseq in more accurately detecting cellular state changes and transcriptional regulation (see methods)^39^. 24h after psilocybin, we dissected mPFC, dissociated, captured and sequenced whole cells and retained 39 435 high-quality cells from 8 independent biological replicates (Fig.1a,b; Extended Data Fig. 1; Tables S2,S3). We identified 11 clusters representing all major cell-types and many neuron subtypes (Fig. 2b), all of which were present in every sample (Extended Data Fig. 1, Table S3). Marker gene expression confirmed cluster cell-type identity (Fig. 2c; Table S4).

**Figure 2.**
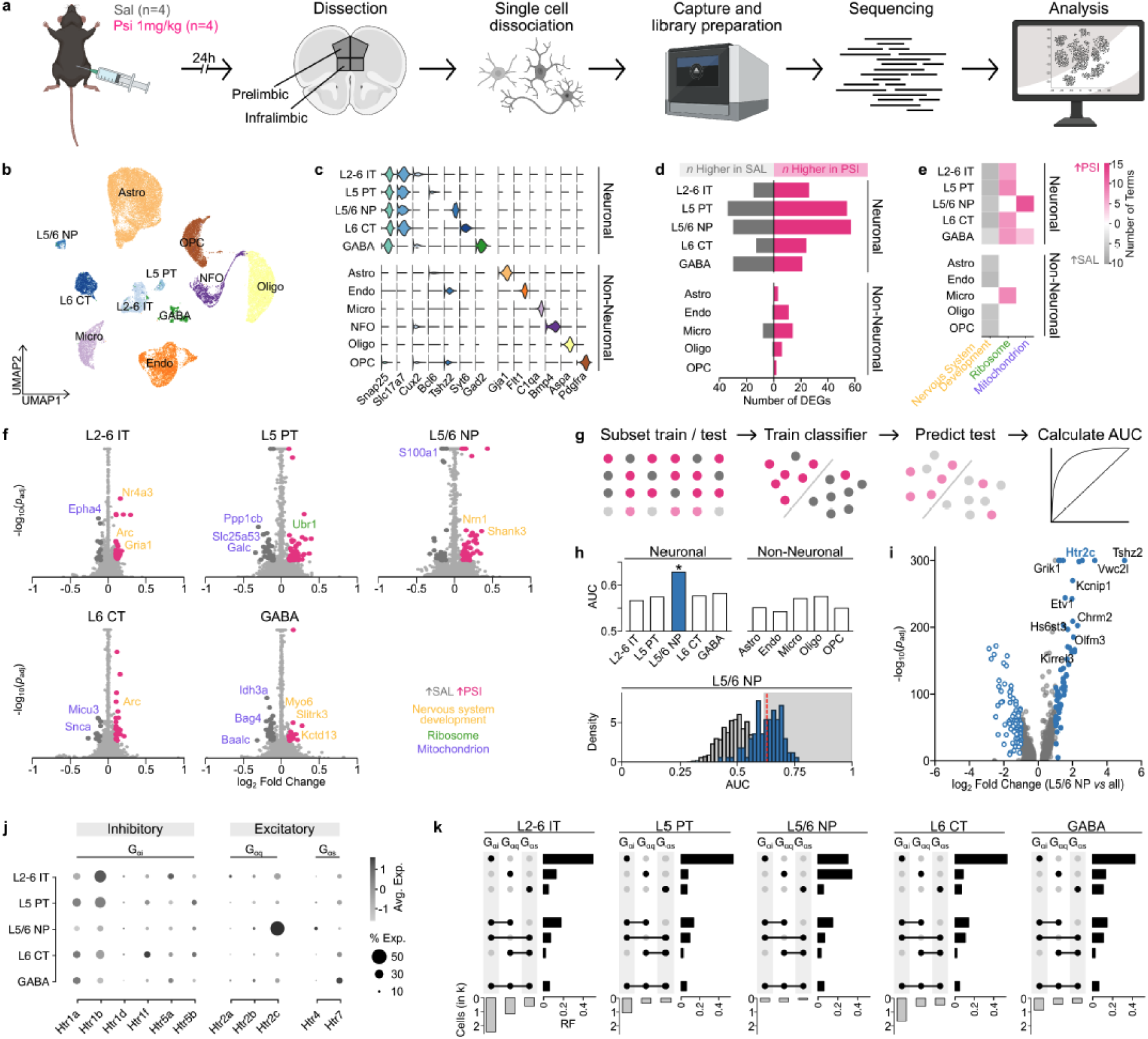
A single injection of psilocybin induces robust transcriptional changes in L5/6 NP neurons. **a.** Schematic of single cell RNA-seq experiment. Mice were injected with psilocybin (1mg/kg; n =4) or saline (n=4) and placed in a clean cage for 30 mins then returned to their home-cage. 24h later infralimbic and prelimbic cortices were dissected and dissociated single-cells were captured for library preparation and sequencing. **b.** UMAP plot to visualize clustering of mPFC neuronal and non-neuronal cell types. **c.** Expression of cell type-specific marker genes in neuronal and non-neuronal cell clusters confirms cell-type identities. **d.** Bar plot showing the total number of genes up (pink) or downregulated (grey) by psilocybin in each cell-type cluster. The largest number of differentially expressed genes were identified in neurons and specifically in L5/6 NP and L5 PT neurons. Significance determined by absolute Log2FC>0.1 and FDR corrected p-value <0.05. **e.** Heatmap illustrating the total number of Gene Ontology terms enriched in genes up (pink) or downregulated (grey) by psilocybin in each neuronal and non-neuronal subtype. GO terms are grouped by their ancestor terms to allow comparison across cell-types. Up-regulated genes in L5/6 NP neurons predominantly enrich for *nervous system development* terms whereas in other excitatory neurons up-regulated genes enrich for *ribosome* terms. *Mitochondrion* terms are enriched in downregulated genes across cell types. **f.** Volcano plots depict genes significantly up (pink) or downregulated (grey). Gene name color denotes associated gene ontology (orange, *nervous system development*; blue, *mitochondrion*; green, *ribosome*).**g.** Schematic of AUGUR cell type prioritization for quantifying transcriptional impact of psilocybin across cell types.^42^ Briefly, AUGUR trains a classifier to predict group (saline, psilocybin) in each cell type using a subset of cells, then estimates classifier performance in held out cells to derive the area under the curve (AUC) to quantify the relative impact of psilocybin in each cell type. **h.** Bar plot of AUGUR classifier performance (area under the ROC curve) for each neuronal and non-neuronal cell type. Larger AUC indicates superior classifier performance, and so greater impact of psilocybin on transcription in this cell type. Permutation testing confirmed L5/6 NP predicts treatment effect above chance (*p<0.05) and this exceeds all other cell types (One-way ANOVA, F=9.873, p= 5.81x 10^-16^, post-hoc Tukey’s Range Test, max p.adj= 0.0004; grey area is significantly higher than permutation distribution; red dotted line depicts L5/6 NP AUC mean). **i.** Volcano plot visualizing genes that distinguish L5/6 NP from other excitatory neurons. *FindAllMarkers* in Seurat compared gene expression in L5/6 NP neurons to all other excitatory neurons. Positive Log2 Fold Change indicates genes with higher expression in L5/6 NP. **j.** Dot plot depicts serotonin receptor expression in mPFC neuron subtypes measured in a spatial transcriptomic dataset^44^. Receptors are segregated by canonical G-protein coupling (G_Ds_, G_Dq and_ G_Di_) the predominant associated effect of G-protein signaling (i.e. excitatory, inhibitory). **k.** UpSet plot summarizes patterns of cell-type specific gene expression and co-expression of 5-HTR genes observed at the level of individual single cells. In most neuron subtypes, the largest number of cells exclusively express transcripts for 5-HTRs that couple to G_Di_ (e.g. *Htr1a*, *Htr1b*). Uniquely in L5/6 NP neurons, the largest number of cells exclusively express transcripts for 5-HTRs that couple to G_Dq_ (e.g. *Htr2c*). OPC: oligodendrocyte precursor cell; Oligo: oligodendrocyte; NFO: newly formed oligodendrocyte; Micro: microglia; Endo: endothelial cell; Astro: astrocyte; GABA: GABAergic neurons; L6CT: Layer 6 corticothalamic neurons; L5/6 NP: Layer 5/6 near projecting neurons; L5 PT: Layer 5 pyramidal tract neurons; L2-6 IT: Layer 2-6 intratelencephalic neurons. B-F, bars represent mean +/- SEM * p<0.05, **p<0.01,***p<0.001. Graphics were made using BioRender.

To understand how psilocybin induces lasting changes in mPFC, we probed psilocybin-induced gene expression changes in each cell type. Using a generalized linear mixed model^40^ to optimize statistical power while controlling for non-independence of variability (i.e. cells nested within replicates), we identified hundreds of genes that were differentially expressed (DEGs) between saline and psilocybin groups across cell-types (Fig. 2d; Table S5,S6). Neurons were the primary locus of psilocybin’s sustained effect with the largest number of DEGs. L5/6 NP neurons contained the largest number of upregulated DEGs (57) followed by L5 PT neurons (54) with fewer downregulated DEGs in both L5/6 NP (30) and L5 PT (34). All excitatory neuron subtypes contained more upregulated than downregulated DEGs. GABAergic neurons were unique in having relatively more downregulated DEGs (21 up, 30 down).

To interrogate the biological significance of the sustained transcriptional impact of psilocybin, we used gene set enrichment analysis with clusterProfiler^41^, to identify gene ontologies enriched in genes up or downregulated by psilocybin (Table S5-S8). Distilling the resulting large numbers of enriched gene ontologies to their ancestral terms (Fig. 2e,f; Table S9) revealed that, broadly, psilocybin induced expression of genes associated with nervous system development processes, including regulation of neuron projection development, regulation of vasculature development, regulation of trans-synaptic signaling, and myelination in L5/6 NP neurons (e.g. *Shank3*, *Nrn1*) and, to a lesser degree, inhibitory neurons (e.g. *Myo6*, *Kctd13*). Psilocybin also increased ribosome biogenesis across excitatory and inhibitory neurons (e.g. *Ubr1* in L5 PT, *Ola1* in L6 CT), and broadly suppressed mitochondrial genes across most cell types. Notably, psilocybin also robustly increased expression of immediate early genes (IEGs) in L2-6 IT (e.g. *Arc*, *Nr4a3*) and L6 CT (e.g. *Arc*) and L2-6IT ( e.g. *Nr4a3*). The upregulation of IEGs and synaptic genes suggests sustained increases in activity in these cell-types consistent with increased neuroplasticity and altered network activity.

### Cell-type prioritization confirms psilocybin robustly regulates L5/6 NP neurons

Binding and receptor expression profiles predict that psilocybin could exert broad effects. Single-cell sequencing can simultaneously profile all cell types to determine if psilocybin’s effects in mPFC are cell-type specific. To quantitatively assess the impact of psilocybin in each cell type we used AUGUR^42^ for cell type prioritization (Fig. 2g). AUGUR builds a classifier for each cell type from a fixed number of cells to predict the treatment condition and then assesses classifier performance on held out cells, with higher scores (area under ROC curves) indicating stronger impact of psilocybin in this cell type. Comparing treatment response across cell types revealed a robust response in L5/6 NP neurons, with above chance prediction that exceeded all other cell types (Fig. 2h; Table S10).

We then asked why this specific excitatory neuron subtype is preferentially responsive to psilocybin. Given the widely held hypothesis that psilocybin preferentially targets specific cell-types based on 5-HT_2A_ expression^30^, we first considered expression of the gene encoding this receptor, *Htr2a* but found only sparse expression in L5/6 NP neurons with higher expression in L2-6 IT neurons and endothelial cells (Extended Data Fig. 5). Overall, we observed relatively modest *Htr2a* expression consistent with reports of lower expression in mPFC than neighboring premotor and motor cortices^43^. To probe the molecular identity of L5/6 NP neurons we compared gene expression in L5/6 NP neurons to all other excitatory neurons in saline samples and identified 67 genes enriched in L5/6 NP neurons. *Htr2c*, the gene encoding the 5-HT_2C_ receptor, was among the top three most highly enriched genes. (Fig. 2i;Table S11). We hypothesized that, distinct from the acute hallucinogenic response, sensitivity to psilocybin’s sustained effects may be conferred by patterns of 5-HT receptors beyond the 5-HT_2A_ receptor.

### L5/6 NP neurons express distinct complements of 5-HT receptors

Individual cells often co-express two or more 5-HTRs such that examining any single subtype in isolation misses the complexity of serotonergic signaling at the cell. This is particularly important given that distinct 5-HTR subtypes couple to different G proteins to mediate sometimes opposing cellular effects. We therefore examined both expression and co-expression of 5-HTR transcripts for receptors in each cell-type. Technical dropouts in scRNA-seq data make zero values difficult to interpret, so to accurately define co-expression patterns, we used a published spatial transcriptomic dataset ^44^. Mapping expression of 5-HTRs across neuronal subtypes in infralimbic and prelimbic cortices confirmed that L5/6 NP neurons display uniquely high expression of *Htr2c* and relatively low expression of all other *Htr* transcripts (Fig. 2j). In contrast, *Htr2c* expression is low in all other neuronal subtypes. Notably, this also confirmed overall low *Htr2a* expression in these cortical subregions.

We then considered the effect of 5-HTRs in each neuronal subtype according to their canonical coupling to G_i_ (*Htr1a, Htr1b, Htr1d, Htr1f, Htr5a, Htr5b*), G_s_ (*Htr4*, *Htr7)* or G_q_ (*Htr2a*, *Htr2b*, *Htr2c*) ^45^. Grouping *Htr* transcripts into subtypes, we plotted the number of cells expressing *only* G_i_, G_s_ or G_q_ associated transcripts as well as those expressing *multiple* classes^14,46^. This revealed that L5/6 NP neurons are indeed unique in that a higher fraction of cells express *only* G_q_ *Htr* transcripts compared to cells *also or only* expressing G_i_ *Htr* transcripts (Fig. 2k). An opposing pattern characterized all other excitatory neuron subtypes, with a higher fraction of cells expressing *only* G_i_ *Htr* transcripts. Overall, the fraction of cells expressing multiple classes of *Htr* transcripts was a minority across all neurons. The observation that L5/6 NP have a high proportion of uniquely G_q_ *Htr* expressing cells aligns with the robust effect of psilocybin in L5/6 NP neurons, suggesting the balance specific receptors may confer cell-type specific sensitivity. This predicts psilocybin will differentially impact neuron subtypes, potentially exerting a stronger excitatory modulation of L5/6 NP neurons and more inhibition of other excitatory neurons.

GABAergic neurons display a mixed *Htr* expression profile predicting mixed effects in inhibitory neurons (Extended Data Fig. 5). Further resolving GABAergic subtypes revealed that heterogeneous *Htr* expression patterns map to specific subtypes with uniquely G_q_ *Htr*-expressing cells predominating in *Sncg^+^* interneurons and uniquely G_i_ *Htr*-expressing cells predominating in all other interneuron subtypes with a sizeable proportion of *Vip^+^* interneurons also expressing either G_q_ or G_s_. This suggests psilocybin will exert highly cell-type specific effects in interneuron subtypes that through integration with distinct interneuron functions and circuit architecture^47^ are poised to exert important effects on network-level neuroplasticity.

### Multivariate analysis identifies transcriptional signatures of psilocybin that associate with a diversity of 5-HTR gene expression across cell types

Differential expression analyses identified transcripts that were up- or downregulated 24h after psilocybin primarily in neurons with fewer changes in non-neuronal cell-types. DEG analysis identifies gene by gene transcript changes that are consistent across a population of cells with each gene considered largely in isolation, overlooking the modular nature of biological gene regulation in that functionally related groups of genes are co-regulated^53^. However, the heterogeneity of 5-HTR expression within and across cell types predicts that psilocybin will exert multi-dimensional effects that DEG will fail to identify. We therefore turned to a multivariate supervised latent factor decomposition approach, partial least squares discriminant analysis (PLS-DA)^54,55^ to identify groups of genes or ‘*modules’* that distinguish between gene expression profiles of psilocybin- and saline-treated mice.

We trained a separate classifier model on gene expression data for each cell type to identify gene modules that predict the effect of psilocybin in each cell-type. We used permutation testing to establish module robustness by comparing the actual explained variance to the null distribution (Fig. 3a). This approach identified multiple robust gene modules in every cell-type, each of which captures unique aspects of the transcriptome-wide effects of psilocybin (Extended Data Fig. 6). This confirms that psilocybin induces sustained effects on transcription in non-neuronal as well as neuronal cells. Comparing the discriminative capacity of the first gene module, which captures the principal axis of treatment-relevant variation, revealed that L5 PT (with an explained variance of 0.60) and L5/6 NP (with an explained variance of 0.57) neurons had the highest discriminative capacity among all cell-types (Fig. 3b). Thus, while psilocybin indeed induces transcriptional alterations across cell-types, the most profoundly impacted cells are two excitatory neuron subtypes, L5/6 NP and L5 PT.

**Figure 3.**
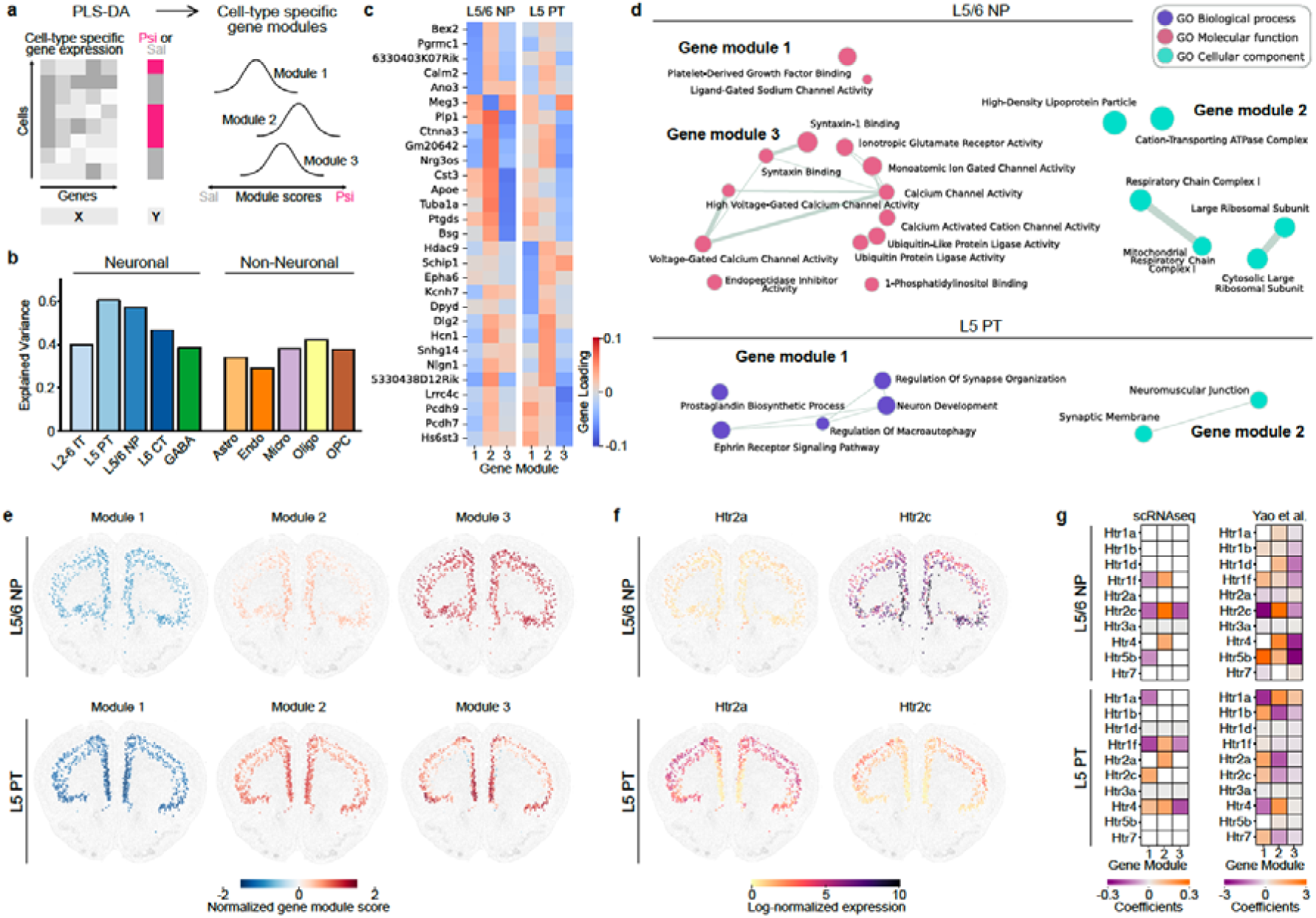
Supervised latent factor modeling reveals cell-type-specific psilocybin sensitivity beyond 5-Htr2A signaling in deep-layer cortical neurons. **a.** Schematic of supervised latent factor decomposition for identifying orthogonal gene modules that discriminate between gene expression profiles from psilocybin- and saline-treated mice, independently within each cell-type. We trained partial least squares discriminant analysis (PLS-DA) models separately for each cell-type to distinguish between cells from saline- and psilocybin-treated mice. Each cell-type model estimates gene expression profiles from all samples and derives a set of latent discriminative modules capturing treatment-related variation. The module scores quantify the extent to which a single cell expresses the psilocybin-associated gene signature captured by a given cell-type-specific module. **b.** Among the eleven cell types analyzed, L5/6 NP and L5 PT neurons exhibited the strongest discriminative capacity, with explained variances of 0.57 and 0.60, respectively. **c.** Heatmap visualizes the contribution of the top 5 genes in each of the three gene modules in L5/6 NP and L5 PT neurons. Each gene module captures a unique pattern of coordinated gene contributions, where positive weights (red) indicate that higher transcript levels, considering co-expression relationships, vary in the same direction as psilocybin treatment, whereas negative weights (blue) reflect variation associated with the saline condition. Note that genes can participate in multiple modules reflecting the multifunctionality of genes. We identified a number of genes associated with neuronal activity and synaptic plasticity including *Meg3*, an activity-induced lncRNA that regulates synaptic plasticity by modulating AMPAR surface expression. **d.** L5/6 NP and L5 PT neurons also showed distinct transcriptional enrichments in individual gene modules, based on Gene Ontology (GO) annotations. All enriched pathways (FDR q-val<0.05) are provided in Supplementary Tables S32-S35. **e.** Normalized expression of the three L5/6 NP and L5 PT gene modules for each cell type projected onto a spatial transcriptomics atlas^64^ from gene module embeddings. Red indicates expression profiles that are more psilocybin-like, and blue, more saline-like. Gene module 1 in each cell type more closely resembles those of saline-treated mice, as expected given that the spatial data were derived from untreated control brains. **f.** Normalized expression of *Htr2a* and *Htr2c* in L5/6 NP and L5 PT neurons. L5 PT cells highly express *Htr2a* whereas *Htr2c* is the primary serotonin receptor expressed in L5/6 NP cells in the spatial atlas. **g.** Regressing *Htr* expression on gene module scores in our scRNAseq dataset and using the spatial atlas as a validation dataset revealed that, in both data sets, expression of *Htr2c*, but not *Htr2a*, is predicted by the transcriptional effect of psilocybin in L5/6 NP cells across all gene modules.

We then looked further into the leading genes in each module. Each gene module captures the unique associations amongst a group of genes, the expression patterns of which predict the effect of psilocybin treatment in that cell-type. Notably, PLS-DA allows a gene to participate in more than one module, reflecting the biological reality of multi-functionality of individual genes, with functions variably defined by the global co-ordination amongst interacting partners^56^. Visualizing the expression weights of the genes that are the top contributors to the module prediction identified many genes associated with neuronal modulation and synaptic plasticity (Fig. 3c). *Meg 3,* an activity-induced lncRNA modulates activity-dependent AMPAR surface expression to regulate synaptic plasticity, is implicated across modules in both L5/6 NP and L5 PT. This gene is particularly interesting given the kind of enduring effects of psilocybin on synaptic transmission we observe in mPFC (Fig. 1d-f) and that others have previously demonstrated in other brain regions^18,19,57^. *Calm2* encodes a calcium binding protein also implicated in regulating synaptic transmission while *Schip1* and *Epha6* are implicated in structural plasticity, and *Ano3* and *Kcnh7* in excitability^58–62^. The epigenetic regulator, *Hdac9,* encodes a histone deacetylase implicated in psychiatric disorders and functional and structural plasticity^63^. Of note, some genes exhibit opposite effects depending on their co-expression partners, as reflected by the presence of both positive and negative loadings for a single gene across different modules, emphasizing the multi-dimensionality of the effects of psilocybin.

We then used gene set enrichment analysis (GSEA) to interrogate the biological, molecular, and cellular pathways psilocybin modulated in each cell-type, here focusing on the cell-types with the largest discriminative capacity (see Tables S32-S35 for full results). This revealed distinct transcriptional programs in individual gene modules in L5 PT and L5/6 NP based on Gene Ontology (GO) annotations (Fig. 3d). L5/6 NP module 1 was uniquely enriched for ‘Ligand-Gated Sodium Channel Activity (GO:0015280)’ and ‘Platelet-Derived Growth Factor Binding (GO:0048407)’ on the GO molecular function atlas. These pathways were exclusive to L5/6 NP cells and not detected among the top three modules of any other cell type. L5/6 NP module 2 was enriched for oxidative phosphorylation pathways, particularly complex I of the electron transport chain as reflected on the GO cellular component atlas. L5/6 NP module 3 strongly enriched for synaptic and ion channel functions. L5 PT module 1 significantly enriched for five pathways in the GO biological processes atlas, including synaptic and neuronal functions. Likewise, L5 PT module 2 enriched for cellular component terms related to synaptic structure.

Having identified robust psilocybin gene modules across cell-types we then asked if these cell-type specific transcriptional programs might relate to *Htr* expression profiles. We first performed this analysis in our scRNA-seq data and then used an independent spatial transcriptomics dataset for validation^64^. To this end, we projected the gene module embeddings onto a spatial transcriptomics atlas to simultaneously visualize expression of serotonin receptors and gene modules on a 2D brain slice^64^.

We aligned and sign-corrected gene modules (PLS components) such that positive scores correspond to gene expression patterns associated with psilocybin treatment, while negative scores reflect patterns more consistent with the saline condition (cf. Methods). In both L5/6 NP and L5 PT neuronal types, the first module—which captures the primary axis of treatment-related transcriptomic variation—exhibited negative scores in the spatial atlas (Fig. 3e). This indicates that the transcriptional profiles in these cell-types in this data set more closely resemble those of saline-treated mice, as expected given that these data are from drug naïve mice. Next, we mapped 5-HTR genes in this spatial dataset and again confirmed that overall *Htr2a* expression is low in the IL/PL sub-regions, although higher in L5 PT than L5/6 NP, and that *Htr2c* is the dominant 5-HTR transcript expressed in L5/6 NP cells (Fig. 3f). We then asked if expression of 5-HTR genes is predicted by gene module expression, first in our scRNA-seq data set and then sought replication in the spatial dataset. Across both datasets, *Htr2c* expression robustly associated with psilocybin treatment in all L5/6 NP modules and in module 1 of L5 PT (Fig. 3g). Interestingly, L5/6 NP modules 1 and 3 have a negative regression coefficient suggesting that the sustained transcriptional effect of psilocybin 24h post injection captured by these modules is *inversely* related to *Htr2c* expression. Given that *Htr2c* expression levels are high in L5/6 NP neurons at baseline, this negative association may suggest a 5-HT_2C_ receptor downregulation following acute psilocybin exposure. Looking across additional cell-types revealed a number of other replicable receptor-gene module predictions including in L2-6 IT neurons where several 5-HTR genes (*Htr1a*, *Htr1b*, *Htr2a*, *Htr2c*, *Htr1f*, *Htr7*) robustly associate with gene module 1, suggesting that psilocybin mediates sustained transcriptional alterations across cell types via diverse 5-HT receptors (Extended Data Fig. 7).

We identified gene module associations with a range of *Htr* genes, consistent with recent findings that at least some sustained effects of psilocybin do not require 5-HT_2A_ receptor signaling^18,19^. In addition to potential serotonin receptor-mediated effects beyond 5-HT_2A_, recent reports suggest psilocin may directly interact with TrkB receptors as well as the Arylhydrocarbon receptor to modulate brain derived neurotrophic factor (BDNF) signaling^28,49^. We therefore asked if expression of *Ntrk2*, the gene encoding TrkB receptors, or *Ahr*, the gene encoding Arylhydrocarbon receptor, might also be tracked by certain gene module expression (Extended Data Fig. 8). Strikingly, in L5 PT neurons, the primary axis of variance (gene module 1) robustly predicted across datasets both *Ntrk2* and *Ahr* expression. Notably, neither *Ahr* nor *Ntrk2* was reliably predicted by gene module expression in L5/6 NP neurons. This further reinforces the observation that while both are robustly regulated by psilocybin, the mechanisms by which this occurs in L5 PT and L5/6 NP neurons are likely distinct. However, *Ntrk2* expression was consistently predicted by the primary axis of variance in GABA neurons and a number of non-neuronal cell types (endothelial, microglia, oligodendrocytes) while *Ahr* expression was tracked by other neuron types (L2-6 IT neurons, L6CT neurons). This adds further weight to experimental observations that signaling via TrkB and AhR receptors mediate some sustained effects of psilocin.

Taken together, we find that psilocybin mediates diverse transcriptional programs across neuronal and non-neuronal cells that associate with a diversity of 5-HTR genes, as well genes for BDNF-signaling related receptors, TrkB and AhR. Overall, the effect of psilocybin is most prominent in L5/6 NP and L5 PT neurons, and in L5/6 NP neurons, transcriptional signatures of psilocybin robustly associate with *Htr2c* expression patterns. This suggests that 5-HT_2C_ signaling plays an important role in the sustained effects of psilocybin.

### 5-HT2c signaling contributes to the neuroplastic effects of psilocybin

Finally, to test the prediction that 5-HT2_c_ signaling contributes to psilocybin’s sustained neuroplastic effects, we injected mice with a specific 5-HT_2C_ antagonist (SB-242084 or vehicle) prior psilocybin injection (1mg/kg) and then prepared acute slices for mEPSC recordings 24h later (Fig. 4a). Relative to neurons from vehicle treated mice, SB reduced average mEPSC amplitude in neurons in L5/6 with a more modest reduction in frequency and a significant shift in the cumulative probability of both amplitude and inter-event-intervals (Fig. 4b-f). To test if SB might simply suppress mEPSC amplitude, we repeated the experiment injecting mice with SB or vehicle followed by only a saline injection. In saline treated mice, SB did not alter frequency or amplitude (Extended Data Fig. 9). Overall, this confirms that 5-HT_2C_ contributes to the sustained neuroplastic effects of psilocybin in the mPFC.

**Figure 4.**
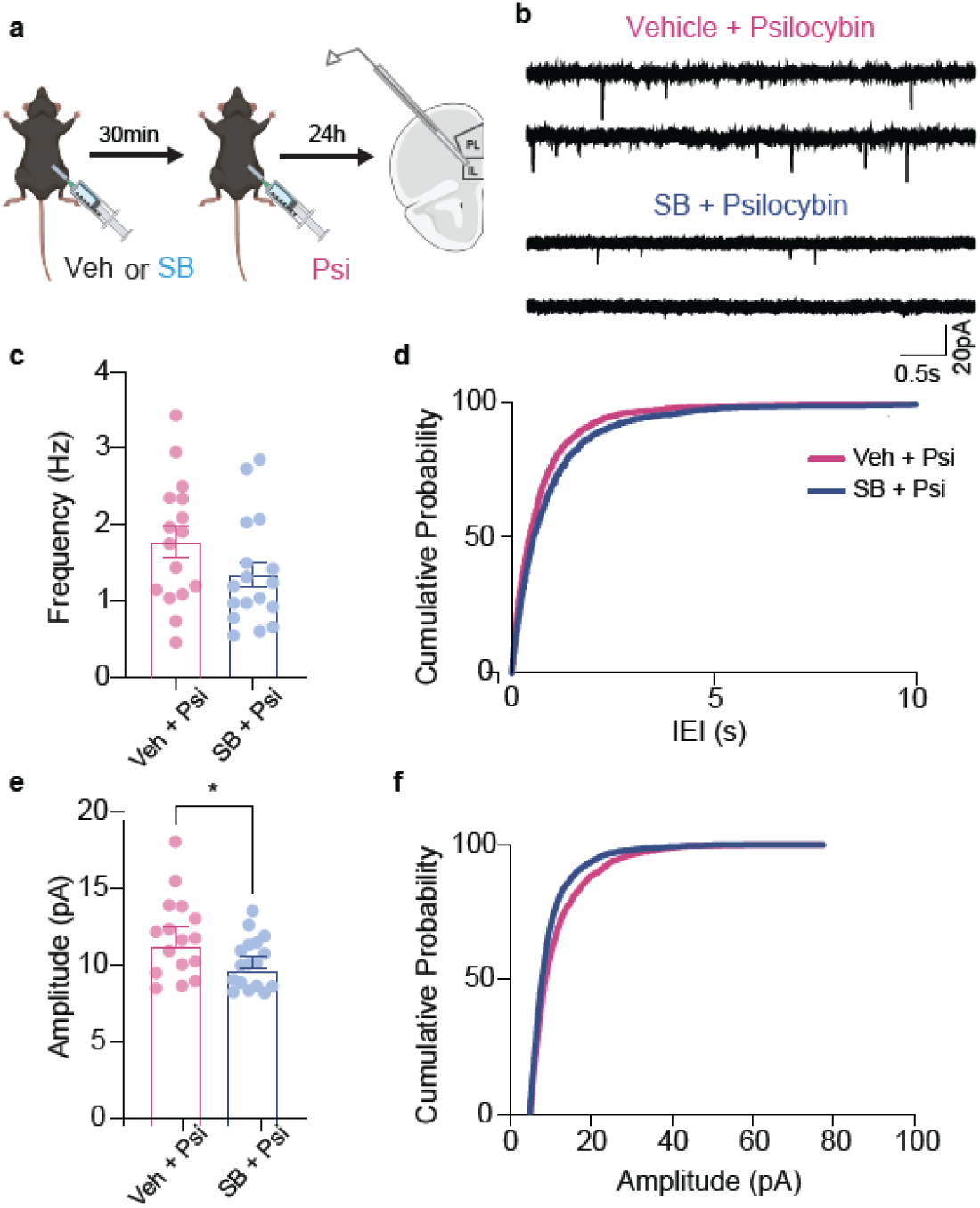
5-HT_2C_ antagonism attenuates effect of psilocybin on synaptic function **a.** Schematic of synaptic physiology experiment. Mice were injected with SB-2242084 (SB) or vehicle and placed in a clean cage for 30 mins then injected with psilocybin (1mg/kg) and returned to the same cage for a further 30 min then returned to their home-cage. 24h later brain slices containing the infralimbic cortex were prepared for patch-clamp recordings. **b.** Representative traces of mEPSC recorded from L5/6 neurons in the infralimbic cortex in brain slices from mice that received SB or vehicle prior to psilocybin 24h before recording. **c.** Bar graphs of mEPSC frequency. Pre-treatment with SB did not significantly alter mEPSC frequency. Each circle denotes a cell (n=21 cells from five mice for SB, and n=16 cells from four mice for vehicle). **d.** Cumulative distribution plots of mEPSC inter-event interval (IEI) for the same recordings. SB decreased IEI. **e,f** Similar to **c,d** but for amplitude. Pre-treatment with SB reduced mEPSC amplitude (p<0.05).

## Discussion

Acute hallucinogenic effects of psilocybin are mediated by the 5-HT_2A_ receptor yet whether this same mechanism also mediates the sustained neuroplastic effects is unclear. Here we show that a single dose of psilocybin increases synaptic transmission in mPFC 24h later and this is accompanied by transcriptomic changes across neuronal and non-neuronal cell types that are inconsistent with an exclusively 5-HT_2A_ mechanism. Psilocybin preferentially impacts L5/6 NP neurons, a type of deep layer excitatory neuron with low expression of the 5-HT_2A_ gene, and high expression of the 5-HT_2C_ gene. Further, *Htr2c* expression is robustly predicted by the expression of psilocybin-associated gene signatures in L5/6 NP neurons, and pharmacologically blocking 5-HT_2C_ before psilocybin injection attenuates the sustained increase in synaptic transmission. These findings establish a key role for 5-HT_2C_ signaling in psilocybin’s neuroplastic effects.

Examining DEGs, cell-type prioritization and multivariate gene module analysis all converged in identifying L5/6 NP neurons as robustly impacted by psilocybin. These deep layer neurons make up approximately 6% of prefrontal cortical neurons and are distinguished from other deep layer neurons by their short-range projections, molecular identity and modular distribution pattern^44^. Like L5 PT neurons, L5/6 NP neurons enrich for *Fezf2*, but are differentiated by uniquely high *Tshz2* expression.

Developmentally, L5/6 NP neurons migrate from the cortical plate earlier than either PT or IT neurons, playing a key role in initial cortical circuit wiring^65^. This relatively small population of excitatory neurons has been largely overlooked until now and there are no validated genetic tools for targeting L5/6 NP neurons. Only by taking an unbiased approach probing all cell types were we able to identify this intriguing modulation by psilocybin. Of note, genetic strategies for isolating L5 PT neurons may also capture L5/6 NP neurons given shared expression of *Fezf2*^30^. Our multivariate analysis confirmed previous findings in other prefrontal cortical areas showing that L5 PT are also highly impacted by psilocybin^30^. Nevertheless, our analyses indicate that, while psilocybin impacts both L5/6 NP and L5 PT, these cell-types express distinct complements of 5-HTRs and psilocybin induces distinct gene expression programs such that the mechanisms and effects of psilocybin in each cell-type are distinct and warrant more granular investigation. As advances in genetic tools push back the boundaries of cell-type specific targeting, it will be exciting to learn more about the functional role of L5/6 NP neurons.

Here, we establish that L5/6 NP neurons are enriched for 5-HT_2C_ receptors and further, that *Htr2c* expression is robustly predicted by psilocybin’s transcriptional effects in L5/6 NP neurons. This informed the hypothesis that 5-HT_2C_ signaling is necessary for psilocybin’s sustained neuroplastic effects. In support of this, pre-treatment with a selective 5-HT_2C_ antagonist attenuated increased synaptic transmission after psilocybin. While much research centers on 5-HT_2A_, psilocin ubiquitously binds 5-HTRs and increasing evidence challenges the exclusive role of 5-HT_2A_ in the sustained neuroplastic effects^18,19,28,66^. The present findings add further weight to the importance of mechanisms beyond the 5-HT_2A_ receptor.

Interrogating single-cell expression patterns of all expressed 5-HTR genes revealed that L5/6 NP neurons are unique in predominantly expressing only G_Dq_-coupled 5-HTR genes, while other excitatory neuron subtypes predominantly express G_Di_-coupled 5-HTR genes. These distinct patterns of 5-HTRs predicts differentially effects of psilocybin, favoring increased L5/6 NP neuron activity, while likely inhibiting other excitatory neurons, potentially accounting for the prominent changes in gene expression in L5/6 NP neurons. Single-cell *Htr* gene expression patterns in GABAergic subtypes revealed that *Sst+*, *Pvalb+*, and Lamp5+ neurons predominantly express genes encoding G_Di_-coupled 5-HTRs, predicting primarily inhibitory effects, whereas *Sncg+* and to some degree *Vip+* interneurons predominantly express genes encoding G_Dq_ - or G_Ds_ - coupled 5-HTRs, predicting primarily excitatory effects. *In vivo*, *Pvalb^+^* and *Sst^+^* monosynaptically inhibit excitatory neurons, while *Vip^+^* interneurons monosynaptically inhibit *Sst^+^ and Pvalb^+^* interneurons to ultimately inhibit the inhibition of excitatory neurons such that psilocybin overall may decrease GABAergic inhibition of excitatory neurons which could contribute to observed increases in excitatory synaptic transmission^47^.

DEG analysis also revealed upregulation of synaptic regulatory genes in GABAergic neurons consistent with suppressed GABAergic function. In GABAergic neurons, psilocybin upregulated *Myo6*, implicated in AMPA receptor endocytosis and weakening of glutamatergic transmission as well as *Kctd13,* which is involved in regulating synaptic transmission and RhoA signaling^67^. Psilocybin suppressed Slitrk3, implicated in GABAergic synapse organization^68^. In excitatory neurons, psilocybin-induced transcriptional programs are consistent with sustained effects of on synaptic function and structure^19^. In L5/6 NP, psilocybin upregulated *Shank3*, a synaptic scaffold implicated in autism, schizophrenia and anxiety, also implicated in antidepressant response^69–71^ as well as *Nrn1*, an activity synaptic plasticity regulator also implicated in antidepressant response^72^. Psilocybin upregulated several activity associated genes including *Arc*, an IEG implicated in synaptic plasticity and antidepressant response, in both L2-6 IT and L6 CT^73^. In L2-6IT, *Nr4a3*, an IEG implicated in synaptic plasticity and learning, was most strongly upregulated by psilocybin^74^. Both psilocybin and ketamine were reported to acutely induce IEG expression in various brain regions^24^. Here we identified sustained activation 24h later, long after psilocybin was fully metabolized, consistent with sustained changes in mPFC activity.

Psilocybin also increased expression of *Gria1*, the key AMPA receptor subunit for activity-dependent plasticity, consistent with established effects on synaptic structure and function and synaptic transmission in mPFC^19,29,75^. *Gria1* upregulation was most prominent in L2-6IT neurons with more marginal effects in L5 PT (*log2FC =* 0.09, *padj* = 0.02) and L5/6 NP (log2FC= 0.14, *p.adj* =0.1) neurons, suggesting effects may be limited to a subset of neurons, paralleling the sparse and heterogeneous maintenance of psilocybin-induced spines^19^. In addition to relatively restricted regulation of synaptic plasticity genes, psilocybin broadly suppressed mitochondrial genes across neuronal and non-neuronal cells. A scRNA-Seq study of stress susceptibility linked high stress vulnerability to both elevated mitochondrial and reduced synaptic gene expression in mouse mPFC^76^. Our findings suggest that psilocybin may induce gene expression programs that oppose certain depression-associated transcriptomic signatures.

Ultimately, we find that psilocybin mediates multi-dimensional transcriptional regulation within and across cell-types. While DEG analyses primarily identified effects in neurons, multivariate analyses revealed psilocybin-associated gene modules across neuronal and non-neuronal cell-types, with the most prominent effects in the same cell-types most enriched for DEGs i.e. L5/6 NP and L5 PT excitatory neurons. Multivariate analyses thus more sensitively captured psilocybin-induced transcriptional modulation while ultimately converging on similar biology. A unique strength of the multivariate approach is in parsing the multiple, distinct patterns of gene expression induced by psilocybin. We also introduce a powerful method for untangling complex, multi-dimensional mechanisms by predicting receptor expression from gene module scores in a manner that is broadly generalizable to any range of potential transcriptional mediators. This method revealed that in L5/6 NP neurons, *Htr2c* expression was robustly predicted by all gene modules, consistent with enrichment of 5-HT_2C_ in this cell-type. Across neuronal and non-neuronal cell-types diverse *Htr* transcripts were predicted by a variety of gene modules, illustrating that psilocybin signals via many 5-HTRs. We also applied this approach to interrogate the role of other receptors previously implicated in psilocybin’s sustained effects, namely the BDNF TrkB receptor, encoded by *Ntrk2,* and ArylHydrocarbon receptor, encoded by *Ahr* ^28,49^. Both, *Ntrk2* and *Ahr* expression were predicted by gene modules in L5 PT, but not L5/6 NP, reinforcing that, while psilocybin prominently impacts both cell-types, the primary mechanisms are likely distinct.

Here we show that a single injection of psilocybin increased synaptic transmission in mPFC 24h later and induced parallel cell-type specific transcriptional alterations that are most prominent in L5/6 NP and L5 PT neurons. We identify a number of molecular mediators of psilocybin’s sustained neuroplastic effects and establish a key role for 5-HT_2C_ receptor signaling. These findings contribute important new insight into how psilocybin interacts with mPFC cell-types. Understanding these cell-type specific changes defines an essential foundation for rigorously understanding how psilocybin induces changes in neural circuit function to mediate the sustained changes in behavior and cognition that underlie psilocybin’s therapeutic efficacy in psychiatric disorders.

## Methods

### Behavioral Methods

#### Animals

7-week-old male and female C57BL/6J mice were obtained from Jackson Laboratories and habituated to the colony room for one week prior to the start of manipulations. Animals were maintained on a 12h light-dark cycle (lights on at 7:00am) at 22-25°C group-housed with same-sex cage-mates (5 mice per cage) with *ad libitum* access to food and water. All experimental manipulations occurred during the light cycle, in accordance with guidelines of McGill University’s Comparative Medicine and Animal Resources Center and approved by the McGill Animal Care Committee.

#### Psilocybin injection

Psilocybin was obtained from the Usona Institute in Fitchburg, Wisconsin under the Investigational Drug & Material Supply Program. A stock solution was prepared by dissolving psilocybin powder in sterile saline at dilutions of 0.025mg/ml, 0.05mg/ml, 0.1mg/ml and 0.2mg/ml solutions for intraperitoneal injection at 0.01ml/g body weight to probe head-twitch response.

#### Head twitch response

Immediately following intraperitoneal injection of vehicle or drug (0.25, 0.5, 1 or 2mg/kg), animals were placed in a clean cage in a behavioral testing room and video recorded under red light for 30 minutes. Video files were then manually analyzed for the number of head twitches exhibited in a 30-minute testing time frame.

### Ex vivo electrophysiological analysis of synaptic function

#### Psilocybin, saline and SB-242084 injection

To examine the delayed effects of psilocybin, mice were injected with either 0.9% sterile saline (0.01ml/g) or 0.01g/ml psilocybin (1mg/kg) via intraperitoneal injection. After injection, mice were housed individually in a clean cage with a handful of their own home-cage bedding for 30min before being returned to their home cage with cage mates. To examine the effect of blocking the 5-HT_2C_ receptor prior to psilocybin injection, a separate group of mice were pretreated with the 5-HT_2C_ receptor antagonist SB-242084 (SB; Tocris) at a dose of 1 mg/kg. A stock solution of SB was prepared by dissolving it in a vehicle (containing: 5% DMSO, 40% PEG, 5% Tween 80, and 50% sterile water) at a concentration of 0.01 g/ml. Mice received SB (or vehicle) via intraperitoneal injection 30 minutes before psilocybin administration. After the SB injection, mice were housed individually in a clean cage with a handful of their own home-cage bedding until the psilocybin(1mg/kg; or saline) injection. Thirty minutes after psilocybin injection, mice were returned to their home cage with their cage mates for 24 hours.

#### Brain slice preparation

24h after drug injection, mice were deeply anesthetized. Transcardial perfusion was performed with 20-25 ml of ice-cooled carbogenated NMDG artificial cerebrospinal fluid (aCSF): (containing in mM: 92 NMDG, 2.5 KCl, 1.25 NaH_2_PO_4_, 30 NaHCO_3_, 20 HEPES, 25 glucose, 2 thiourea, 5 Na-ascorbate, 3 Na-pyruvate, 0.5 CaCl_2_·4H_2_O and 10 MgSO_4_·7H_2_O. Osmolarity: ∼310 mOsmol/L; pH: 7.3–7.4). Brain slices (200 μm) were prepared using a vibratome (Leica VT1200S) in ice-cooled carbogenated NMDG aCSF using. All brain slices recovered in 32–34 °C carbogenated NMDG aCSF for 10 min. Then, they were transferred into room-temperature carbogenated HEPES holding aCSF (containing in mM: 92 NaCl, 2.5 KCl, 1.25 NaH_2_PO_4_, 30 NaHCO_3_, 20 HEPES, 25 glucose, 2 thiourea, 5 Na-ascorbate, 3 Na-pyruvate, 2 CaCl_2_·4H_2_O and 2 MgSO_4_·7H_2_O. Osmolarity: ∼310 mOsmol/L; pH: 7.3–7.4). Brain slices were kept in HEPES holding aCSF until recording.

#### Voltage-clamp recording

Recordings targeting infralimbic cortex were performed in room-temperature carbogenated aCSF (containing in mM: 128 NaCl, 3 KCl, 1.25 NaH_2_PO_4_, 2 MgCl_2_, 2 CaCl_2_, 24 NaHCO_3_ and 10 glucose. Osmolarity: ∼310 Osm; pH 7.2). All signals were amplified and digitized by Multiclamp 700B (Molecular Device) and Digidata 1550B (Molecular Device) respectively. Traces were filtered with a 2 kHz Bessel low-pass filter and data were acquired at 10 kHz. Pipette electrodes (2.5 to 5 MΩ) were filled with a cesium-methanesulfonate based internal solution (containing in mM: 130 Cs-methanesulfonate, 10 HEPES, 0.5 EGTA, 8 NaCl, 5 TEA-Cl, 4 Mg-ATP, 0.4 Na-GTP, 10 Na-phosphocreatine and 1 QX-314. Osmolarity: 290-300 mOsmol/L; pH 7.2). Neurons were held at a membrane potential of –70 mV during recordings. Miniature Excitatory Postsynaptic Currents (mEPSCs) were recorded in the present of picrotoxin (50 μM, Sigma) and TTX (0.5 μM, Hello Bio).

Cells with access resistance above 20 MΩ or with access resistance increased by > 20% were rejected. For each brain slice, at least one neuron from L2/3 and one from L5/6 was recorded to examine the effect of psilocybin. For the slices used to examine the effect of blocking the 5-HT_2C_ receptor, only neurons from L5/6 were recorded. Miniature data analysis was conducted offline using Mini Analysis (6.0.3, Synaptosoft, USA).

### Single Cell RNA-seq Sample Preparation and Sequencing

#### Considerations in method selection

While technically demanding in brain tissue, scRNAseq offers important advantages over single nucleus RNAseq (snRNAseq). Because snRNAseq is limited to nuclear transcripts, far fewer RNAs are captured, and gene length biases and lack of cytoplasmic RNA species limit inference about RNA in the whole cell^77^. While both methods are powerful in identifying cellular heterogeneity, by capturing cytoplasmic and nuclear RNA, scRNAseq more accurately detects cellular state changes and differential transcriptional regulation^39^. Notably, snRNAseq experiments generally need to sequence many more nuclei than is the case cells in scRNAseq experiments to offset the fewer transcripts that are captured by snRNAseq.

#### scRNAseq tissue preparation

A separate cohort of female mice were decapitated 24h post saline or psilocybin injection. Brain tissue was dissociated as described in^78^. Brains were extracted, immediately submerged in ice-cold Hibernate-A / B27 / Glutamax medium (HABG; 60 ml of Hibernate A medium with 1 ml of B27 and 0.15 ml of GlutaMAX) and subsequently sliced into 1 mm coronal slices using a cooled brain matrix. Bilateral infralimbic and prelimbic cortices were microdissected in an agarose Petri dish with 2 ml of HABG. Next, the dissected tissue was chopped into small pieces of approximately 0.5 x 0.5 x 0.5 cm and transferred into 5 ml of HABG supplemented with 45 μM actinomycin. After equilibrating the tissue to 30°C for 8 minutes in a water bath, the tissue was transferred into calcium-free Hibernate-A supplemented with 2 mg ml−1 of papain, 45 μM actinomycin, and 2x GlutaMAX for enzymatic dissociation at 30°C for 30-35 minutes in a shaking water bath (170 rpm).

Following dissociation, the tissue was transferred into 2 ml HABG supplemented with 45 μM actinomycin and equilibrated to RT for 5 minutes. Using fire-polished Pasteur glass pipettes, samples were next mechanically dissociated by triturating suspensions approximately 10 times in 45 seconds. Upon allowing the remaining tissue pieces to settle, the supernatant was passed through a 30 μm filter to remove large debris and cell aggregates. This step was repeated 2 times, until all tissue pieces were dissociated.

To remove any remaining debris, the resulting 6 ml supernatant was carefully layered onto an OptiPrep gradient as described in^79^. Briefly, this gradient sorts cell material into debris and 4 fractions: fraction 1 enriches for oligodendrocytes; fraction 2 enriches for debris; fraction 3 enriches for neurons; fraction 4 enriches for microglia.

Following a 15 minute centrifugation (800 g, 4°C), we discarded the debris, fraction 1 and fraction 2; collected fraction 3 in a new tube; and discarded fraction 4. Fraction 3 was then diluted with 5 ml HABG and centrifuged for 3 minutes (200g, 4°C). The supernatant was discarded, and the pellet was resuspended in 5 ml of DPBS supplemented with 0.04% BSA. After a final centrifugation for 3 minutes (200g, 4°C), the supernatant was discarded and the final pellet was resuspended in 50 ul of DPBS supplemented with 0.04% BSA.

#### scRNA-seq Library Preparation

Single-cell suspensions were washed and resuspended in PBS with 0.04% BSA. We aimed to capture ∼8,000 cells per sample. An aliquot of cells was used for LIVE/DEAD viability testing (Thermo Fisher Scientific). Single-cell libraries were generated using the 10x Genomics Chromium Controller or Chromium X instrument and Chromium Next GEM Single Cell 3[ GEM, Library & Gel Bead Kit v3.1 (10x Genomics) according to the manufacturer’s protocol. Briefly, cells suspended in reverse transcription reagents, along with gel beads, were segregated into aqueous nanoliter-scale gel bead-in-emulsions (GEMs). The GEMs were then reverse transcribed in a T1000 Thermal cycler (Bio-Rad) programed at 53°C for 45 min, 85°C for 5 min, and hold at 4°C. After reverse transcription, single-cell droplets were broken and the single-strand cDNA was isolated and cleaned with Cleanup Mix containing DynaBeads (Thermo Fisher Scientific). cDNA was then amplified with a T1000 Thermal cycler programed at 98°C for 3 min, 12 cycles of (98°C for 15 s, 63°C for 20 s, 72°C for 1 min), 72°C for 1 min, and hold at 4°C. Subsequently, the amplified cDNA was fragmented, end-repaired, A-tailed and index adaptor ligated, with SPRIselect Reagent Kit (Beckman Coulter) with cleanup in between steps. Post-ligation product was amplified with a T1000 Thermal cycler programed at 98°C for 45 s, 12 cycles of (98°C for 20 s, 54°C for 30 s, 72°C for 20 s), 72°C for 1 min, and hold at 4°C. The sequencing-ready libraries were cleaned up with SPRIselect, quality controlled for size distribution and yield (LabChip GX Perkin Elmer), and quantified using qPCR (KAPA Biosystems Library Quantification Kit for Illumina platforms). Libraries were loaded on MGI DNBSEQ-G400 and sequenced using the following parameters: 28 bp Read1, 8 bp Index i7, and 150 bp Read2. Libraries were sequenced at an average depth of 40,000 reads/cell.

### scRNA-seq Data Analysis

#### Pre-processing and quality control

CellRanger v7.0.0 was used with option “count” and default parameters to filter and align raw reads to the pre-compiled mouse reference refdata-gex-mm10-2020-A. CellBender v0.3.0 was then used to remove background RNA^80^. Further quality control and processing were performed with Seurat v4.4.0^81^. Cells were filtered on the number of genes (greater than 800 and within two standard deviations from average), the number of total reads (within two standard deviations from average) and mitochondrial content (lower than 10%). In each sample, only genes detected in more than 3 cells were retained. After quality control, two saline-injected samples were pooled due to low cell number.

#### Identification of cell types

Cell type-specific markers were identified by detecting differentially expressed genes between the given cell type and the remaining cells using *FindAllMarkers* function of Seurat^81^ package with default parameters. Excitatory neuron sub-type specific markers were identified by detecting differentially expressed genes between the given subtype and the remaining excitatory neurons, thereby excluding markers shared across all excitatory neuron subtypes. SciBet^82^, SingleCellNet^83^ and SingleR^84^ were used to annotate cell types with two published datasets as references. The first dataset consists of cells collected from saline treated P60 animals published by Bhattacherjee, et al^85^. The second dataset consists of cells from the Allen Mouse Brain Atlas with region label ILA;PL;ORB (infralimbic, prelimbic and orbitofrontal areas)^86^. A consensus annotation was assigned when at least two methods agreed. Cells were only assigned as neuron if labeled as neuron by both references. Neuron subtypes were labeled from the Allen Mouse Brain Atlas. The expression of cell type and neuron subtype markers was examined to confirm the cell type assignment.

#### Normalization, Dimension Reduction and Integration of Samples

The dataset was scaled to 10,000 reads per cell and log transformed. Samples were integrated by merging samples without batch correction. Variable features were identified with the variance-stabilizing transform method. Dimension reduction was performed with the 2000 most variable features and the top 30 dimensions were used for clustering and projection to two dimensions with Uniform Manifold Approximation and Projection.

#### Differential Expression Analysis

Differential gene expression analysis was performed for each individual cell type with Libra using a negative binomial generalized linear mixed model to account for nesting of cells within biological replicates with offset and Wald test^40^. Genes with absolute Log2 Fold Change greater than 0.1 and adjusted p-value (Benjamini-Hochberg FDR correction) less than 0.05 were considered to be differentially expressed.

#### Gene Set Enrichment Analysis

Gene symbols were first converted to Entrez ID using the DAVID conversion tool (https://david.ncifcrf.gov/helps/conversion.html). Genes without converted Entrez ID were discarded. GSE analysis was then performed using clusterProfiler^87^. To compare ontologies across cell-types, the ancestors of the enriched Gene Ontology terms for each cell-type were obtained using EMBL-EBI’s QuickGO REST API (https://www.ebi.ac.uk/QuickGO/api/index.html) and aggregated for the top 50 of either positively or negatively enriched terms in each cell type or subtype.

#### Cell Prioritization

Augur^42^ was used to probe the magnitude of response of each cell type/subtype to psilocybin injection. AUGUR is optimized for statistical inference at the level of cell types such as quantitatively evaluating the involvement of specific cell types in response to a therapeutic intervention by using a machine-learning framework to rank cell types according to the relative magnitude of response to a treatment. The larger the response to treatment in any given cell type, the greater the separability of cells by experimental group. Briefly, AUGUR trained a machine-learning classifier to predict the experimental condition (saline or psilocybin) associated with each cell.

Separate classifiers were trained for each cell-type, and accuracy evaluated by cross-validation. The accuracy measure quantifies the magnitude of transcriptional response to treatment in each cell type. A parameter sweep was performed to determine the best combination. The combination of vquant=0.9, ntree=150, mtry=20 optimized prediction and was used for further analysis. The cell condition labels were then shuffled (with Augur “permute” option and otherwise identical parameters as abovementioned) to determine whether the Augur score was significantly above chance.

#### Identification of Genes Enriched in L5/6 NP

To identify genes enriched in L5/6 NP neurons separate to any effects of psilocybin, L5/6 NP enriched genes were identified using only cells from saline samples. The *FindAllMarkers* function of the Seurat package was used with default parameters to compare L5/6 NP cells against all remaining excitatory neurons.

#### Serotonin(co-)expression analysis

Spatial transcriptomic MERFISH data^44^ from were used to analyse co-expression of HTR receptors in the PL and IL areas. MERFISH data provides more accurate quantification of reads with fewer technical dropouts compared to single-cell RNA sequencing. The increased coverage of GABAergic cell types in this dataset also allowed for a more detailed investigation of HTR expression in these subtypes. The normalized count matrix was downloaded from^44^. For the main analysis, we subset to 7 HTRs, classified according to canonical G-protein of the associated receptor: G_Di_ (Htr1a, Htr1b, Htr1d, Htr1f, Htr5a, Htr5b), G_Dq_ (Htr2a, Htr2b, Htr2c) and G_Ds_ (Htr4, Htr7) based on literature^46,88^. UpSet plots were generated on all cells that expressed at least one inhibitory or one excitatory receptor using the *ComplexHeatmap* package in R with the ‘distinct’ setting.

#### Supervised latent factor modeling of transcriptome-wide gene modules predictive of psilocybin treatment

We used partial least squares analysis (PLS) to deconvolve and extract high-dimensional latent components—referred to as “gene modules”^55^—from normalized scRNA-seq count matrices obtained from psilocybin- and saline-treated mice. PLS generates distinct, uncorrelated modes of variation: linear combinations of gene expression features in X that are optimally aligned with variation in a target outcome variable Y. Using PLS for the purposes of classification has been referred to as PLS discriminant analysis (PLS-DA)^54^.

We considered a set of 8,460 marker genes from a spatial transcriptomics atlas^64^, of which 8,264 overlapped with our experimental dataset and were retained for inclusion in the PLS-DA model. However, PLS-DA is known to be prone to overfitting when the number of features greatly exceeds the number of observations—a common risk in high-dimensional transcriptomic data, where high collinearity is also prevalent^54^.

In our case, for most cell types, the number of genes (features) exceeded the number of transcriptomic profiles (observations), with the exception of astrocytes and newly formed oligodendrocytes. To mitigate the risk of overfitting, we first applied principal component analysis (PCA) to reduce the dimensionality of the normalized gene expression matrices across all cell types, ensuring consistency in the modeling pipeline. For each cell type, we computed the top 100 components of variation in the data, which were then used as features for the PLS-DA analysis. This preprocessing step has been shown to improve model robustness and performance in PLS-DA applications^56^. Accordingly, in the PLS-DA model, the explanatory input matrix was defined as the top 100 principal components, computed separately for each major brain cell type. The corresponding outcome variable was constructed based on the binary treatment status, with ‘psilocybin’ coded as +1 and ‘saline’ coded as −1^55,89^. For each cell type independently, we considered two corresponding variable sets, X and Y:

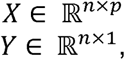

where *n* denotes the number of cell observations (i.e., the top 100 principal component expression vectors from a given cell type) and denotes the number of features (i.e., 8,264 gene expression measurements). PLS-DA finds latent source of variation in and simultaneously uses the emerging latent structure to predict. The two sets and are decomposed as the dot product of two matrices that represent what is often call the model ‘scores’ (), and model ‘loadings’ (), respectively. The latent factor decomposition of the original ambient variable sets is obtained as follows:

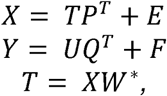

where *T* and *U* are matrices of size *n* x l, *P* is a matrix of size *p* x l, *Q* is a matrix of size *q* x l, and *E* and *F* are matrices of normally distributed error terms for variation not explained in *X* and *Y*, respectively. *l* determines the number of latent variables extracted during the analysis. Following the principle of linear regression, *Y* can be estimated as a function of *X* through the following equation:

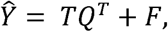

where *T* represents the latent variables of *X*, and *Q* are the corresponding loadings that describe the relationship between *T* and *Y*. This equation can be re-expressed as the multiple regression model:

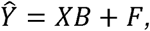

where *B* is a matrix of regression coefficients that maps the predictors in the columns of *X* to the responses *Y*, and *F* represents the residual error matrix. In this form, the relationship between *X* and *Y* is expressed directly, with *B* being derived from the decomposition of *X* and *Y* into their respective latent structures:

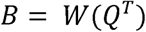

Here, *W* corresponds to the weight matrix associated with *X* and *Q* represents the loadings of the latent structures in *Y*. PLS-DA then identifies a series of l orthogonal latent variables, denoted as *t*_l_, which are obtained through linear combinations of *X* that maximize covariance with Y while remaining uncorrelated with each other. These latent variables are naturally ordered, from highest to lowest, according to the amount of variance in outcome *Y* that they explain. Formally, the optimization problem can be described as follows:

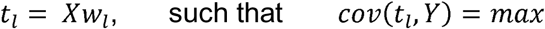

The goal of PLS-DA specification in the context of our study was to independently identify, within each cell type, linearly independent gene modules that capture expression alterations distinguishing the transcriptional profiles of psilocybin- and saline-treated mice. By design, PLS derives orthogonal latent gene modules that capture distinct aspects of psilocybin’s delayed effects, ranked by their ability to explain variance between the transcriptome arrays pertaining to psilocybin- and saline-treated mice. To recover gene-level contribution from the PLS loadings obtained in the PCA-reduced space, we translated them back into the original gene expression space using the PCA rotation matrices, separately for each cell type. This enabled us to express the resulting gene modules on a transcript-by-transcript basis, preserving both biological relevance and interpretability.

To ensure that the gene modules were statistically robust, we performed a rigorous robustness test of each obtained gene module, separately in each cell type. Starting off by considering 20 candidate gene modules in each cell type, we employed a label shuffling permutation test^89,90^ where the binary treatment labels (i.e., 1=‘psilocybin’, −1=‘saline’) were randomly shuffled across cell transcriptome observations, in 100 separate empirical permutation iterations. We then recalculated the otherwise identical cell-type specific PLS-DA models based the permuted outcome data column Y. In each iteration, we computed Pearson’s correlation coefficients between the PLS-DA component between the derived latent embedding space (representing gene module expression) and the (randomly shuffled) outcome label Y across cell observations. The aggregate statistics computed across the 100 iterations were used to construct empirical null distributions for each cell type, modeling the relationship between gene modules and psilocybin treatment under the null hypothesis that no robust gene modules can be extracted that are associated with the treatment in the transcriptomes of the mice under study. To draw inference on gene module stability, we compared the actual achieved explained variance metric (Pearson’s correlation coefficients in the latent embedding space) derived from the original, unshuffled data. In so doing, we quantified how confidently it exceeds the corresponding empirical null distribution – how much above chance level - for each PLS-DA component of a given cell type.

To facilitate module interpretation, we aligned and sign-adjusted the component loadings so that positive scores indeed correspond to gene expression patterns associated with psilocybin treatment, and negative scores correspond to patterns associated with saline treatment. We determined this orientation by calculating the Pearson correlation between the PLS embeddings of X and Y via their derived latent scores and the treatment labels (i.e., 1=‘psilocybin’, −1=‘saline’). If both correlations (X scores vs. labels and Y scores vs. labels) were negative, we flipped the sign of the loadings for that component by multiplying them by −1, ensuring consistent alignment with the treatment labels. This strategy for addressing flipped (“mirrored”) components has been adopted in analogous PLS-DA pipelines^55,56^. This alignment of obtained model parameters was then preserved for all downstream analyses.

Lastly, we performed a one-sample bootstrap hypothesis test to identify, among the 8,264 examined genes, those that robustly contributed to psilocybin treatment detection within each cell-type-specific gene module. Across 1,000 iterations, cells were randomly resampled with replacement from the original cell-type observations prior to applying dimensionality reduction. This approach, previously applied in similar contexts^55,56^, simulates random draws from the broader population of the same cell type. Dimensionality reduction (PCA) and PLS model estimation were then performed on each resampled dataset. To ensure consistency between bootstrap and reference PLS models in each cell-type, we aligned the components from each bootstrap PLS solution to the original PLS solution. Specifically, for each bootstrap replicate, the x-loadings (gene weights) were matched to the reference components by maximizing their Pearson’s correlation. Each matched component was also oriented to have the same sign as the corresponding reference component, ensuring that the directionality of gene contributions was consistent across replicates. This procedure produces an aligned, sign-corrected x-loadings matrix for each bootstrap PLS model, directly comparable to the original PLS solution^56^. For each cell-type-specific module, the top 100 robust genes per module are provided in Supplementary Tables S12–S21, while all genes that passed the bootstrap robustness criterion (2.5–97.5% confidence interval) are listed in Supplementary Tables S22–S31.

#### Gene set annotation analyses

To contextualize our identified psilocybin-predictive gene modules by means of previously established biological processes and pathways, we performed gene set enrichment analysis (GSEA), for all gene modules that have met our robustness criteria (cf. above)^92^. As a preparatory step, we ranked the 8,264 genes in each module based on their PLS-DA module weights, which indicate the strength of contribution of each gene to predicting psilocybin treatment. We then used the *GSEApy* Python implementation (version 1.1.8) of GSEA^93^ with the prerank enrichment tool, which accepts user-provided ranked gene lists. We performed the analysis with the following parameters: *gseapy.prerank* (min_size=5, max_size=8264, permutation_num=1000). We used a fixed random state initialization for reproducibility. We reported the pathways that turned out to be robust according to a FDR threshold of q=0.05. This step was repeated for all gene modules in each of the cell types.

For our reference biological pathway library, we mined i) a general and ii) a mitochondria specific pathway ontology. First, we selected the Gene ontology (GO) knowledgebase, updated as of 2023^94^ to thoroughly evaluate the broad spectrum of biological processes represented by our psilocybin-predictive gene modules. We investigated three levels of pathway annotations: biological process (27,993 annotated terms), molecular function (11,271 annotated terms), and cellular components (4,039 annotated terms). By integrating these reference libraries across different levels of biological information, we aimed to paint a comprehensive portrait of psilocybin-driven transcriptome-wide alterations.

To visualize significant enrichment results (FDR threshold of q = 0.05), we constructed enrichment networks using the enrichment_map function from GSEApy and the NetworkX library (version 3.4.2) for graph generation and layout. For the top-performing cell types—those achieving the highest discriminative capacity between psilocybin- and saline-treated mice—we selected non-zero, significantly enriched gene sets (nz_gsea_df_sig) and computed pairwise similarity metrics (Jaccard and overlap coefficients) based on shared driving genes. These similarity scores defined the edges of the network, with nodes representing individual gene sets. Node size reflected the proportion of input genes contributing to the enrichment signal (Hits_ratio), and edge width was scaled according to the Jaccard coefficient, emphasizing functionally related pathways. Node positions were initialized using a spiral layout to minimize overlap, and labels were repositioned toward the centroid of each node’s neighborhood to improve readability. Nodes were color-coded based on the reference atlas/annotation source associated with each gene set.

#### Mapping gene module expression across the mouse brain spatial transcriptome

We leveraged a comprehensive, high-resolution spatial transcriptomic atlas to map the distribution of our psilocybin-predictive gene modules across the adult mouse brain. This resource provides spatial transcriptomic profiles for over 4.3 million cells^64^, derived from a single male mouse using multiplexed error-robust fluorescence in situ hybridization (MERFISH). Of the 8,460 marker genes included in the imputed version of the atlas, 8,264 overlapped with our experimental dataset and were retained for inclusion in the PLS-DA model (see above).

Among the 59 brain slices in the imputed atlas, we identified slice C57BL6J-638850.56 as the best match to our experimental section. This slice contained 57,175 cells, encompassing 16 major cell classes and 73 subclasses, and was used for all downstream analyses. Note that the scRNA-seq experiment was restricted to IL and PL subregions of mPFC but here we include all cells of a cell-type located within this coronal section irrespective of cortical sub-region. Cells were registered in the Allen Mouse Brain Common Coordinate Framework. We matched our experimentally defined cell types to subclass-level annotations from the reference taxonomy.

Specifically, L5/6 NP was mapped to “032 L5 NP CTX Glut”, L6 CT to “030 L6 CT CTX Glut”, and L5 PT to “022 L5 ET CTX Glut”, and L2-6 IT to “006 L4/5 IT CTX Glut.” The subclass “007 L2/3 IT CTX Glut” was also evaluated as a potential match for L2-6_IT, but it showed a weaker correspondence compared to “006 L4/5 IT CTX Glut” and was therefore not retained for downstream analyses. Non-neuronal populations were also aligned: Oligo to “327 Oligo NN”, OPC and NFO to “326 OPC NN”, Micro to “334 Microglia NN”, Astro to “319 Astro-TE NN”, and Endo to “333 Endo NN”. Inhibitory neurons were grouped into a single GABAergic category, which included the subclasses “052 Pvalb Gaba”, “053 Sst Gaba”, “046 Vip Gaba”, “049 Lamp5 Gaba”, and “047 Sncg Gaba”, as these best matched our experimental annotations. These subclass assignments were used for all downstream analyses.

#### Exploring associations between serotonergic and non-serotonergic receptor expression and psilocybin gene modules expression

We first examined the spatial expression patterns of serotonin receptor genes in the MERFISH atlas. Ten serotonin receptor genes were shared between our experimental dataset and the spatial atlas: *Htr1a, Htr1b, Htr1d, Htr1f, Htr2a, Htr2c, Htr3a, Htr4, Htr5b*, and *Htr7*. Additionally, we included two non-serotonergic receptors, *Ahr* and *Ntrk2*, given that they been recently implicated in mechanisms of psilocybin’s enduring effects on neuroplasticity ^49,95^. This enabled us to quantify, for each cell type independently, the expression levels of individual receptors on the brain slice that best matched our experimental data, while accounting for both the spatial distribution and relative abundance of different cell types.

We then applied the PLS-DA embeddings—previously trained on the corresponding cell types from the psilocybin-treated experimental cohort—to the MERFISH atlas data. This approach allowed us to map the expression of cell-type-specific psilocybin response signatures across the spatial transcriptomic dataset within each matched cell type, thereby assessing psilocybin-associated gene module expression in healthy tissue at high spatial resolution in an independent dataset.

Finally, we evaluated the relationship between cell-type-specific expression of the ten serotonin receptors and the top three psilocybin-related gene modules identified per cell type, which are by construction the modules most explanatory of psilocybin effects, using linear regression. Specifically, for each receptor–module pair, we regressed serotonin receptor expression on gene module scores, resulting in 10 × 3 receptor–module regression models per cell type. Analogous regression models were built for Ahr and Ntrk2. These analyses enabled us to quantify, independently within each cell type, both the extent and identity of receptors whose expression is reflected by transcriptome-wide variation associated with the delayed response to psilocybin.

## Data & Code Availability

The RNA sequencing data generated in this study have been deposited in the NCBI Gene Expression Omnibus (GEO) repository under accession number GSE283929. The data will be released upon publication of this manuscript. A secure token has been created to allow review of record GSE283929 while it remains in private status: mjyviuuuvhgvzqh. All code used to analyze and visualize the single cell RNA-seq data with pre-processed data can be found in the github repo https://github.com/DelongZHOU/Psilocybin.Female.Mouse.mPFC. The repo will be made public when the manuscript is published.

## Acknowledgements

Psilocybin was obtained through the Usona Institute’s Investigational Drug & Material Supply Program. Usona Institute had no involvement in the study design, funding, execution or interpretation. Some figure elements were made using BioRender. We thank Juan-Pablo Lopez for advice on implementing single-cell preparations in adult brain. We thank Claudia Kleinman for advice on bioinformatic analyses. This work was supported by funding from CQDM, HBHL and the Ludmer Centre for Neuroinformatics and Mental Health. We thank the McGill Genome Centre for technical support.

## Author contributions

R.C.B. conceived of the study. H.S., V.C., D.Z and A.S.H. conducted scRNAseq experiments. H.S., D.Z. and C.S analyzed scRNAseq data. Y.C.T. conducted electrophysiological recordings. H.S. analyzed MERFISH data. J.M. conducted and analyzed head twitch experiments. J.R. oversaw scRNAseq capture and sequencing. R.C.B., H.S., D.Z., V.C., C.S. and D.B. interpreted the results and wrote the manuscript.

