## Supplementary File for "Cell-type specific transcriptional modulation by psilocybin induces sustained plasticity in mouse medial prefrontal cortex"

**Extended Data Figures**

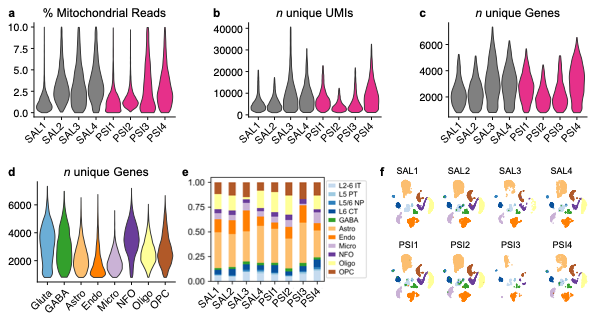

**Extended Data Figure 1. Single Cell RNA-Seq quality control metrics confirm consistent high quality across replicates and groups**

**a.** Percent of mitochondrial read count in each replicate from psilocybin treated (PSI) and saline treated (SAL) mice. **b.** Number of unique UMIs in each replicate from psilocybin treated (PSI) and saline treated (SAL) mice. **c.** Number of unique genes per replicate from psilocybin treated (PSI) and saline treated (SAL) mice. **d.** Number of unique genes for each cell type. **e.** Cell type composition of each replicate from psilocybin treated (PSI) and saline treated (SAL). **f.** UMAP of individual libraries confirms stable cell type proportions across replicates of psilocybin (PSI) and saline treated (SAL) mice. Colors correspond to cell-type markers in (**e**). Gluta: glutamatergic neurons; L2-6 IT: layer 2-6 intratelencephalic neurons; L5/6 NP: layer 5/6 near projecting neurons; L5 PT: layer 5 pyramidal tract neurons; L6 CT: layer 6 corticothalamic neurons; GABAergic neurons; Astro: astrocyte; Endo: endothelial cells; Micro: microglia; NFO: newly formed oligodendrocyte; Oligo: oligodendrocyte; OPC: oligodendrocyte precursor cell

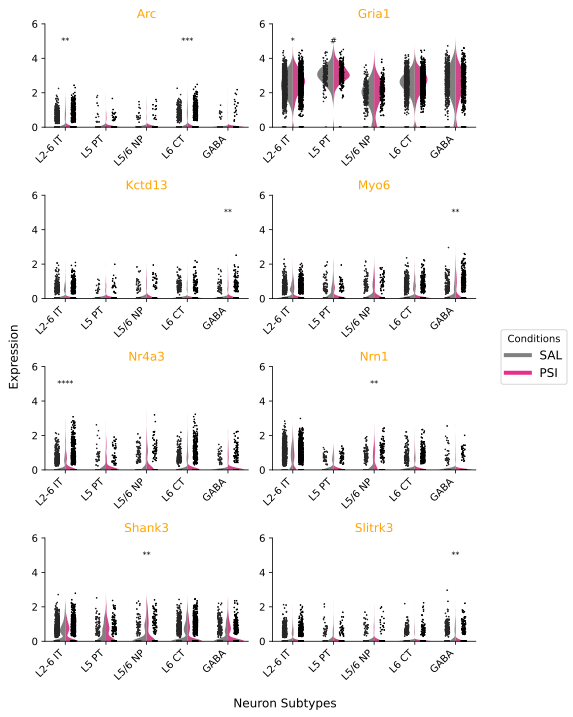

**Extended Data Figure 2. Psilocybin upregulates expression of genes associated with nervous system development gene ontologies** Violin plots show the distribution of expression of genes in individual cells across neuron subtypes in saline (SAL) and psilocybin (PSI) replicates. *p adj.<0.05, log2FC>0.1, ** p adj. <0.01, log2FC>0.1 *** p adj. <0.001, log2FC>0.1 **** p adj. <0.0001, log2FC>0.1, # p<0.05, log2FC<0.1.

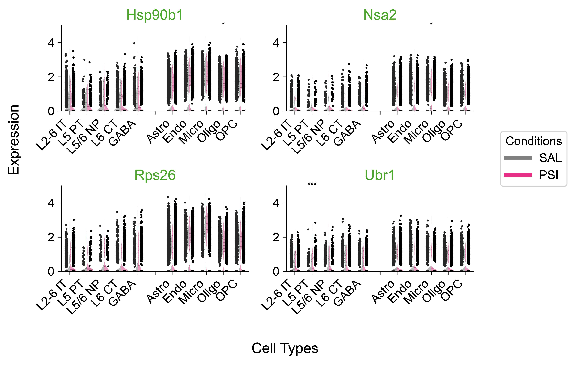

**Extended Data Figure 3. Psilocybin upregulates expression of genes associated with ribosome gene ontologies**

Violin plots show the distribution of expression of genes in individual cells across neuron subtypes in saline (SAL) and psilocybin (PSI) replicates. *p adj.<0.05, log2FC>0.1, ** p adj. <0.01, log2FC>0.1 *** p adj. <0.001, log2FC>0.1 **** p adj. <0.0001, log2FC>0.1

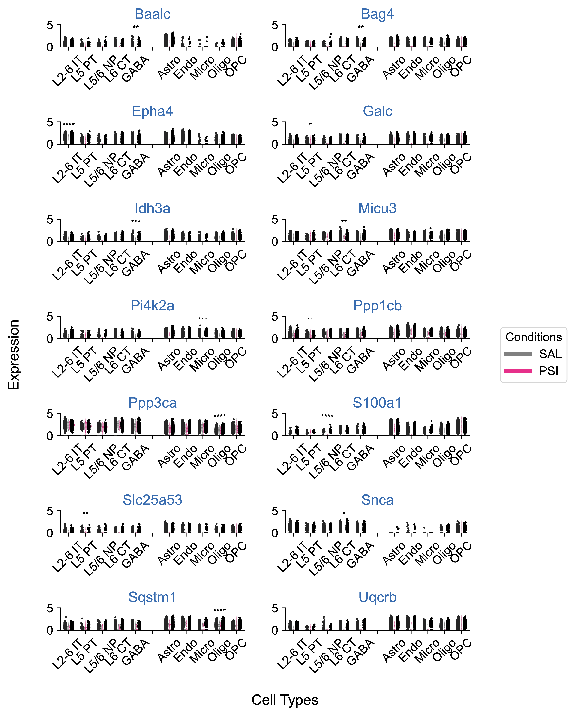

**Extended Data Figure 4. Psilocybin downregulates expression of genes associated with mitochondrial gene ontologies**Violin plots show the distribution of expression of genes in individual cells across neuron subtypes in saline (SAL) and psilocybin (PSI) replicates. *p adj.<0.05, log2FC>0.1, ** p adj. <0.01, log2FC>0.1 *** p adj. <0.001, log2FC>0.1 **** p adj. <0.0001, log2FC>0.1

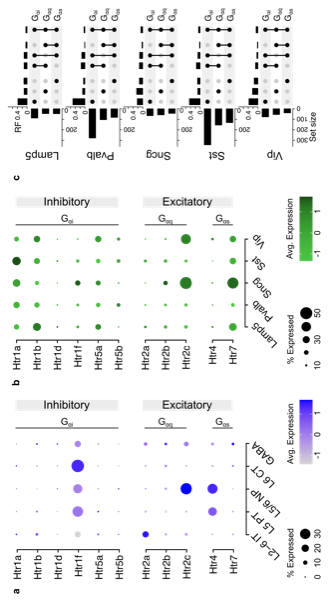

**Extended Data Figure 5. Serotonin receptor gene expression by cell-type in scRNA-seq data and in GABA subtypes**

**a.** Dotplot showing serotonin receptor expression in single cell sequencing dataset. **b.** Dot plot depicts serotonin receptor expressions in mPFC GABAergic interneuron subtypes measured in spatial transcriptomic dataset^1^. Receptors are segregated by canonical G-protein coupling (G_ɑs ,_ G_ɑq and_ G_ɑi_) the predominant associated effect of G-protein signaling (i.e. excitatory, inhibitory). **c.** UpSet plot summarizes patterns of cell-type specific gene expression and co-expression of 5-HTR genes observed at the level of individual single cells for subtypes of GABAergic interneurons.  L2-6 IT: layer 2-6 intratelencephalic neurons; L5/6 NP: layer 5/6 near projecting neurons;  L5 PT: layer 5 pyramidal tract neurons;   L6 CT: layer 6 corticothalamic neurons;  GABAergic neurons; Astro: astrocyte; Endo: endothelial cells;  Micro: microglia;  NFO: newly formed oligodendrocyte;  Oligo: oligodendrocyte;  OPC: oligodendrocyte precursor cell

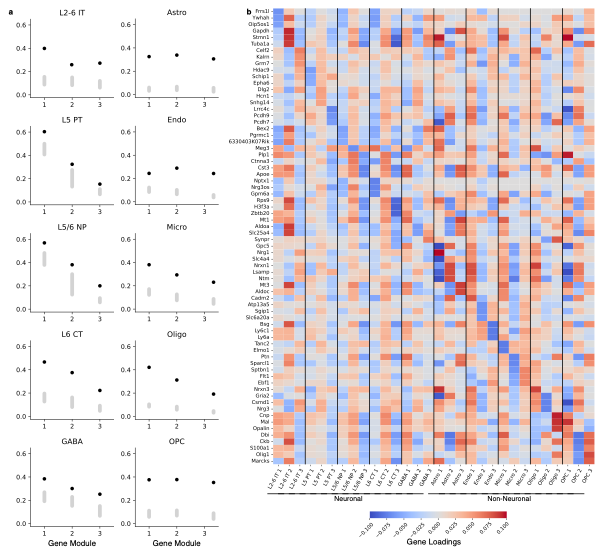

**Extended Data Figure 6. PLS-DA identifies psilocybin-induced transcriptional changes across neuronal and non-neuronal cell types.** We applied a supervised latent factor decomposition approach to identify gene modules that distinguish between psilocybin- and saline-treated mice across ten brain cell types. The top three gene modules for each cell type are shown, collectively capturing distinct aspects of psilocybin's long-term effects. **a.** We assessed the statistical robustness of each gene module within each cell type using a label shuffling permutation test across 100 iterations. We compared the actual explained variance (Pearson’s correlation coefficient, *colored dot*) derived from the original, unshuffled data to the corresponding null distribution (5–95% confidence interval) for each PLS component of a given cell type (*grey bar*). Our analysis revealed the strongest discriminative capacity in L5/6 NP and L5 PT neurons, with explained variances of 0.57 and 0.60, respectively. Among glial cell types, oligodendrocytes ("oligo") showed the highest explained variance (0.42). **b.** Heatmap visualizes the contribution of the top 5 genes in the top three gene module for each cell type. Each gene module captures a unique pattern of coordinated gene contributions, where positive weights (red) indicate that higher transcript levels, considering co-expression relationships, vary in the same direction as psilocybin treatment, whereas negative weights (blue) reflect variation associated with the saline condition. Note that genes can participate in multiple modules reflecting the multifunctionality of genes.

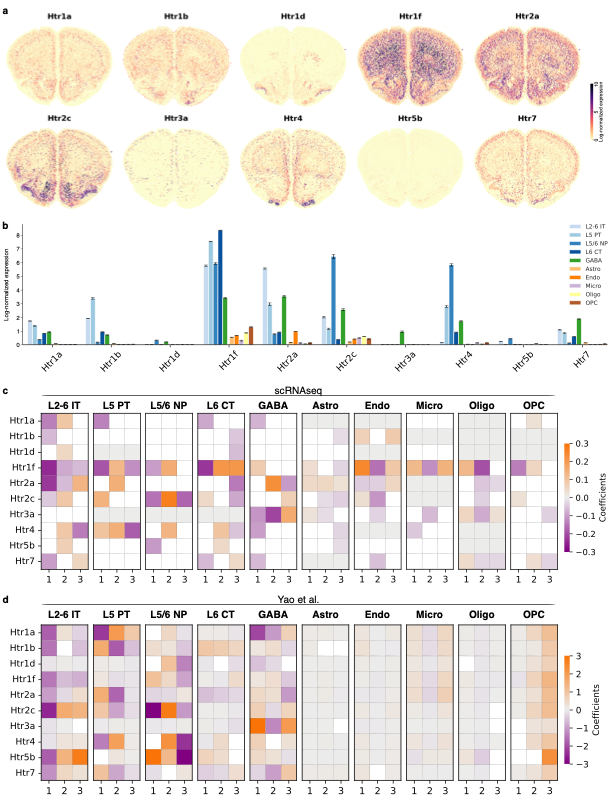

**Extended Data Figure 7. Psilocybin-induced transcriptional changes associate with diverse serotonin receptors across and within cell types.**

**a.** Log normalized expression of the nine serotonin receptors that are present in the scRNAseq dataset across cell types in the spatial atlas^5^ and, **b**., average log normalized expression of these receptors in each cell type accompanied by 95% confidence intervals.  *Htr2a* was most strongly expressed in L2-6 IT neurons (log₁₀(expr) = 5.573 [5.531–5.615]), and other serotonin receptors are also prominent. *Htr1f*  is particularly enriched in L6 CT neurons (log₁₀(expr) = 8.361 [8.347–8.375]), and, again, *Htr2c* is preferentially expressed in L5/6 NP cells (log₁₀(expr) = 6.439 [6.285-6.594]). We evaluated the relationship between cell-type-specific expression of the nine serotonin receptors and the top three psilocybin-related gene modules identified in each cell type using linear regression first in our scRNAseq dataset (**c**) and then validated this in the spatial atlas data set^5^ (**d**). Specifically, for each receptor–module pair, we regressed serotonin receptor expression on gene module scores, yielding 10 × 3 regression models per cell type. This approach allowed us to quantify the extent to which serotonin receptor expression are tracked by the transcriptome-wide variation associated with psilocybin response. Only coefficients that are statistically significant are visualized (α<0.05). While *Htr2a* expression was associated with the psilocybin response in L2-6 IT neurons in both our scRNAseq dataset (β_module 1_=-0.229, *p*<0.001) and the spatial atlas (β_module 1_=-0.992, *p*<0.001)—consistent with its elevated expression in this cell type—we found that other serotonin receptors tracked psilocybin-related transcriptional changes in different cell types. Notably, *Htr2c* expression was strongly associated with gene module expression in L5/6 NP neurons in both our scRNAseq dataset (β_module 1_=-0.153, *p*=0.002; β_module 2_=0.268, *p*<0.001; β_module 3_=-0.172, *p*<0.001) and the spatial atlas (β_module 1_=-2.955, *p*<0.001; β_module 2_=2.788 *p*<0.001; β_module 3_=-1.048, *p*<0.001), reflecting its expression pattern. In contrast, *Htr1f* was linked to psilocybin-responsive modules in L5 PT (β_module 1_=-0.190, *p*<0.001) and L6 CT cells (β_module 1_=-0.232, *p*<0.001) in our scRNAseq dataset, but showed a weaker association in the spatial atlas. Within each cell type, associations tended to be strongest for a specific serotonin receptor; however, individual gene modules often showed significant associations with multiple receptors. These findings suggest that psilocybin-induced transcriptional regulation involves a broad range of serotonin receptors—both across and within cell types—highlighting the need to look beyond the widely studied 5-HT_2A_.

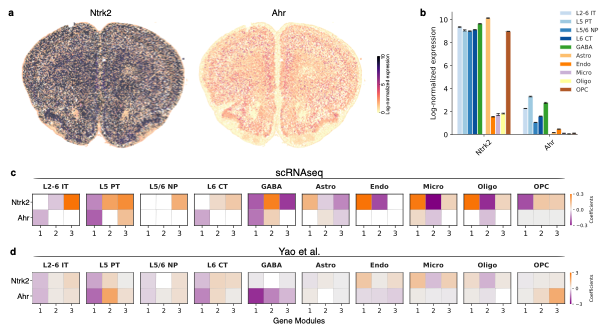

**Extended Data Figure 8. Psilocybin-induced transcriptional changes associate with transcript expression of gene encoding the BDNF receptor, *Ntrk2*, and Arylhydrocarbon receptor, *Ahr* across a range of cell types. a.** Normalized expression of the *Ntrk2* and *Ahr* in the scRNAseq dataset across cell types in the spatial atlas^5^ and, **b**., average expression of these receptors in each cell type accompanied by 95% confidence intervals. We evaluated the relationship between cell-type-specific expression of *Ntrk2* and *Ahr* and the top three psilocybin-related gene modules identified in each cell type using linear regression first in our scRNAseq dataset (**c**) and then validated this in the spatial atlas data set^5^ (**d**). Specifically, for each receptor–module pair, we regressed serotonin receptor expression on gene module scores, yielding 2 × 3 regression models per cell type. This approach allowed us to quantify the extent to which *Ntrk2* and *Ahr* expression is tracked by the transcriptome-wide variation associated with psilocybin response. Only coefficients that are statistically significant are visualized (α<0.05). In L5 PT neurons gene module 1 robustly predicted both *Ntrk2* and *Ahr* expression in our scRNAseq dataset (*Ntrk2*: β_module 1_=-0.127, *p*=0.011; *Ahr:* β_module 1_=-0.176, *p*<0.001) and the spatial atlas (*Ntrk2*: β_module 1_=-0.902, *p*<0.001; *Ahr:* β_module 1_=-1.654, *p*<0.001). Notably, neither *Ahr* nor *Ntrk2* reliably predicted gene module expression in L5/6 NP neurons, again suggesting that while L5 PT and L5/6 NP neuros are both robustly regulated by psilocybin, the mechanisms by which this occurs are likely distinct. *Ntrk2* expression consistently predicted the primary axis of variance in GABA neurons and a number of non-neuronal cell types (endothelial, microglia, oligodendrocytes) while *Ahr* expression was predictive in other neuron types (L2-6 IT neurons, L6CT neurons).

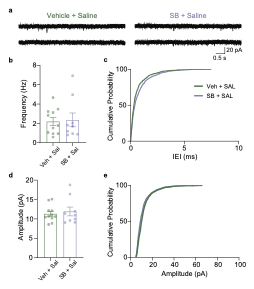

**Extended Data Figure 9. 5HT_2C_ antagonism does not alter synaptic function in vehicle treated mice** Mice were injected with SB-2242084 (SB) or vehicle and placed in a clean cage for 30 mins then injected with saline (1mg/kg) and returned to the same cage for a further 30 min then returned to their home-cage. 24h later brain slices containing the infralimbic cortex were prepared for patch-clamp recordings. **a.** Representative traces of mEPSC recorded from L5/6 neurons in the infralimbic cortex in brain slices from mice that received SB or vehicle prior to saline 24h before recording. **b.** Bar graphs of mEPSC frequency. Pre-treatment with SB did not significantly alter mEPSC frequency. Each circle denotes a cell (n=9 cells from three mice for SB, and n=11 cells from four mice for vehicle). **d.** Cumulative distribution plots of mEPSC inter-event interval (IEI) for the same recordings. **e.** Similar to **b, c** but for amplitude. Pre-treatment with SB did not significantly alter mEPSC amplitude.

**Supplementary Material**

**Supplementary Tables**

**Table S1.** Head twitch responses 30 minutes following 0.25mg.kg, 0.5mg/kg, 1mg/kg or 2mg/kg psilocybin (PSI) or saline (SAL) injection.

**Table S2.** Single-cell library quality control metrics in psilocybin (PSI) and saline (SAL) samples.

**Table S3.** Number of cells for each cell type. L2-6 IT: layer 2-6 intratelencephalic neurons; L5/6 NP: layer 5/6 near projecting neurons; L5 PT: layer 5 pyramidal tract neurons; L6 CT: layer 6 corticothalamic neurons; GABAergic neurons; Astro: astrocyte; Endo: endothelial cells; Micro: microglia; NFO: newly formed oligodendrocyte; Oligo: oligodendrocyte; OPC: oligodendrocyte precursor cell

**Table S4.** Cell-type marker genes enriched in each cell type compared to all other cell types in the full dataset. Gluta: all glutamatergic neurons; L2-6 IT: layer 2-6 intratelencephalic neurons; L5/6 NP: layer 5/6 near projecting neurons; L5 PT: layer 5 pyramidal tract neurons; L6 CT: layer 6 corticothalamic neurons; GABAergic neurons; Astro: astrocyte; Endo: endothelial cells; Micro: microglia; NFO: newly formed oligodendrocyte; Oligo: oligodendrocyte; OPC: oligodendrocyte precursor cell

**Table S5.** List of significantly differentially expressed genes in each cell type between saline (SAL) and psilocybin (PSI) replicates. Significance threshold was set at p adj. <0.05 and log^2^FC>0.1. Gluta: all glutamatergic neurons; L2-6 IT: layer 2-6 intratelencephalic neurons; L5/6 NP: layer 5/6 near projecting neurons; L5 PT: layer 5 pyramidal tract neurons; L6 CT: layer 6 corticothalamic neurons; GABAergic neurons; Astro: astrocyte; Endo: endothelial cells; Micro: microglia; NFO: newly formed oligodendrocyte; Oligo: oligodendrocyte; OPC: oligodendrocyte precursor cell

**Table S6.** Threshold-free differential expression list in each cell type between saline (SAL) and psilocybin (PSI) replicates.

Gluta: all glutamatergic neurons; L2-6 IT: layer 2-6 intratelencephalic neurons; L5/6 NP: layer 5/6 near projecting neurons; L5 PT: layer 5 pyramidal tract neurons; L6 CT: layer 6 corticothalamic neurons; GABAergic neurons; Astro: astrocyte; Endo: endothelial cells; Micro: microglia; NFO: newly formed oligodendrocyte; Oligo: oligodendrocyte; OPC: oligodendrocyte precursor cell

**Table S7.** ID conversion between Gene Symbols and Entrez ID

**Table S8.** Gene Set Enrichment Analysis for all cell types.

Gluta: all glutamatergic neurons; L2-6 IT: layer 2-6 intratelencephalic neurons; L5/6 NP: layer 5/6 near projecting neurons; L5 PT: layer 5 pyramidal tract neurons; L6 CT: layer 6 corticothalamic neurons; GABAergic neurons; Astro: astrocyte; Endo: endothelial cells; Micro: microglia; NFO: newly formed oligodendrocyte; Oligo: oligodendrocyte; OPC: oligodendrocyte precursor cell

**Table S9.** Ancestor terms for Gene Set Enrichment Analysis enriched terms in Table S12.

Gluta: all glutamatergic neurons; L2-6 IT: layer 2-6 intratelencephalic neurons; L5/6 NP: layer 5/6 near projecting neurons; L5 PT: layer 5 pyramidal tract neurons; L6 CT: layer 6 corticothalamic neurons; GABAergic neurons; Astro: astrocyte; Endo: endothelial cells; Micro: microglia; NFO: newly formed oligodendrocyte; Oligo: oligodendrocyte; OPC: oligodendrocyte precursor cell

**Table S10**. AUGUR Cell Prioritization scores of neuron subtypes. Gluta: all glutamatergic neurons; L2-6 IT: layer 2-6 intratelencephalic neurons; L5/6 NP: layer 5/6 near projecting neurons; L5 PT: layer 5 pyramidal tract neurons; L6 CT: layer 6 corticothalamic neurons; GABAergic neurons.

**Table S11.** List of genes enriched in L5/6 NP compared to other excitatory neuron subtypes in saline replicates.

**Tables S12-S21.** Top 100 robust genes in the three psilocybin-associated gene modules in each cell-type.

S12: Astro

S13: Endo

S14: GABA

S15: L2-6 IT

S16: L5/6 NP

S17: L5 PT

S18: L6 CT

S19: Micro

S20: OPC

S21: Oligo

**Tables S22-S31.** All robust genes in the three psilocybin-associated gene modules in each cell-type.

S22: Astro

S23: Endo

S24: GABA

S25: L2-6 IT

S26: L5/6 NP

S27: L5 PT

S28: L6 CT

S29: Micro

S30: OPC

S31: Oligo

**Tables S32- S35:** All enriched pathways (FDR q-val<0.05) identified by gene set enrichment analysis on psilocybin-associated gene modules in each cell-type.

S32: Biological Processes

S33: Cellular Compartment

S34: Molecular Function

S35: MitoCarta
